# Early clonal dominance at priming sets the trajectory for broad HIV serum neutralization

**DOI:** 10.64898/2026.03.04.709621

**Authors:** Bo Liang, Yuxin Zhu, Ryan S. Roark, Xuduo Li, Nitesh Mishra, Christian L. Martella, Anh L. Vo, Gabriella Giese, Qin Huang, Andrea Biju, Lifei Tjio, Rohan Roy Chowdhury, Prabhgun Oberoi, Khaled Amereh, Areeba A. Wani, Yuexiu Zhang, Sharaf Andrabi, Tharunika V. Sekar, Anjali Somanathan, Muzaffer Kassab, Rebecca Nedellec, Sean Callaghan, Gabriel Avillion, Mark G. Lewis, Sara Dutton Sackett, Ashwin N. Skelly, Frederic Bibollet-Ruche, Lawrence Shapiro, Zizhang Sheng, Bryan Briney, Beatrice H. Hahn, Dennis R. Burton, Darrell J. Irvine, Peter D. Kwong, George M. Shaw, Raiees Andrabi

## Abstract

Germline-targeting vaccines aim to elicit HIV-1 broadly neutralizing antibodies (bnAbs), but determinants of serum bnAb induction remain unclear. We previously showed that Q23-APEX-GT2 primes diverse V2-apex bnAb precursors in outbred rhesus macaques (RMs). To assess their maturation into authentic bnAbs, we infected immunized RMs with CAP256.SU SHIV. Four of six developed heterologous serum neutralization breadth, with one neutralizing ∼70% of viruses tested. Early autologous neutralization and B-cell priming efficiency post-prime correlated with post-infection breadth. Longitudinal lineage tracing in lymph-nodes and blood revealed that recruitment and expansion of diverse long-CDRH3 V2-apex B-cell precursors, followed by early dominance of one or two clones, predicted serum breadth. Structural analyses showed that non-neutralizing lineages failed despite cross-reactive Env binding, owing to unfavorable V2-apex approach angles and limited glycan engagement. Together, these findings identify priming efficiency and early clonal dominance as key determinants of serum bnAb induction and provide a framework for HIV vaccine design.

**ONE SENTENCE SUMMARY:** Germline-targeting priming of diverse V2-apex bnAb precursors, coupled with early clonal dominance, determines serum neutralization breadth by enabling selective expansion and maturation of functional lineages into broadly neutralizing antibodies.

## Introduction

The elicitation of broadly neutralizing antibodies (bnAbs) remains a central goal of HIV-1 vaccine development (*1–4*). Although bnAbs arise naturally in a small subset of infected individuals (*5–11*), they typically require years of antigen exposure and extensive affinity maturation to overcome the structural constraints of the heavily glycosylated HIV-1 Envelope (Env) (*7, 12–20*). Germline-targeting strategies aim to initiate bnAb development by engaging rare B cells expressing appropriate antibody precursors (*21–32*). However, successful priming does not necessarily guarantee the development of broad serum neutralization responses (*23–26, 28, 31, 33–37*). The factors governing this transition remain poorly understood, including how early bnAb B cell precursor recruitment, clonal expansion, and antibody maturation determine whether vaccine-induced responses ultimately acquire serum neutralization breadth (*38–42*).

The V2-apex of HIV-1 Env is a vulnerable site targeted by potent bnAbs that possess long complementarity-determining region 3 loops of the heavy chain (CDRH3) and recognize conserved protein and glycan features (*5, 6, 8, 29, 31, 32, 39, 42–56*). We previously developed Q23-APEX-GT2, a germline-targeting immunogen that recruits diverse V2-apex bnAb precursors in outbred rhesus macaques (*32*). Although vaccination elicited cross-reactive monoclonal antibodies, broad serum neutralization remained largely absent after homologous immunization alone. Whether these Q23-APEX-GT2 vaccine-primed precursors could be recalled by subsequent antigenic exposure and mature into authentic bnAbs remained unknown.

Here, we investigated the development of V2-apex serum neutralization breadth following CAP256.SU SHIV infection of Q23-APEX-GT2 trimer immunized macaques. By integrating longitudinal serum neutralization, B cell repertoire tracing, functional characterization of monoclonal antibodies, and cryo-electron microscopy, we linked early priming responses to subsequent neutralization outcomes. Four of six animals developed broad serum neutralization, with early autologous neutralization and the recruitment and expansion of diverse long-CDRH3 lineages associated with subsequent serum neutralization breadth. Longitudinal analyses further revealed that a small number of dominant lineages could drive serum neutralization, whereas other strongly expanded lineages remained non-neutralizing despite extensive maturation. Structural analyses identified differences in angle of approach, glycan engagement, and epitope recognition that distinguished functional bnAbs from non-neutralizing V2-apex lineages. Overall, our findings identify priming efficiency, early clonal dominance and structural constraints on antibody recognition as key determinants of bnAb development, providing a framework for designing HIV vaccines that elicit broad serum neutralization.

## Results

### Q23-APEX-GT2 trimer immunization enables the development of V2-apex serum neutralization breadth following CAP256.SU SHIV infection

We recently showed that Q23-APEX-GT2, an HIV Env V2-apex bnAb site germline-targeting immunogen formulated with the saponin/MPLA nanoparticle adjuvant (SMNP) (*57*) and administered using a slow-delivery, escalating-dose prime regimen (*58, 59*), engages diverse V2-apex bnAb precursors in outbred macaques. Lymph node-derived monoclonal antibodies (mAbs) from some of these animals exhibited cross-neutralizing activity, although heterologous serum neutralization breadth was largely absent, as would be expected after two homologous immunizations with a single germline-targeting immunogen without subsequent heterologous boosting (*32*). We therefore sought to determine whether Q23-APEX-GT2-primed V2-apex precursors could be recalled following heterologous viral exposure and mature into authentic bnAbs capable of mediating broad serum neutralization.

To address this question, we infected Q23-APEX-GT2-immunized macaques with CAP256.SU SHIV at week 18, eight weeks after the second immunization (Fig. 1A) (*38, 60*). CAP256.SU Env was selected because of its propensity to elicit V2-apex bnAbs in both humans and nonhuman primates (*8, 42, 50, 56*). Importantly, this Env preserves key features of the native V2-apex bnAb epitope, including the N156 and N160 glycans and the positively charged strand-C, while maintaining a glycan shield over the V1 and V2 regions that minimizes potential off-target responses to exposed glycan holes (Fig. 1, B to D). This experimental design allowed us to test whether heterologous viral exposure could recall Q23-APEX-GT2 vaccine-primed V2-apex responses and drive their further maturation toward broad serum neutralization.

**Fig. 1.**
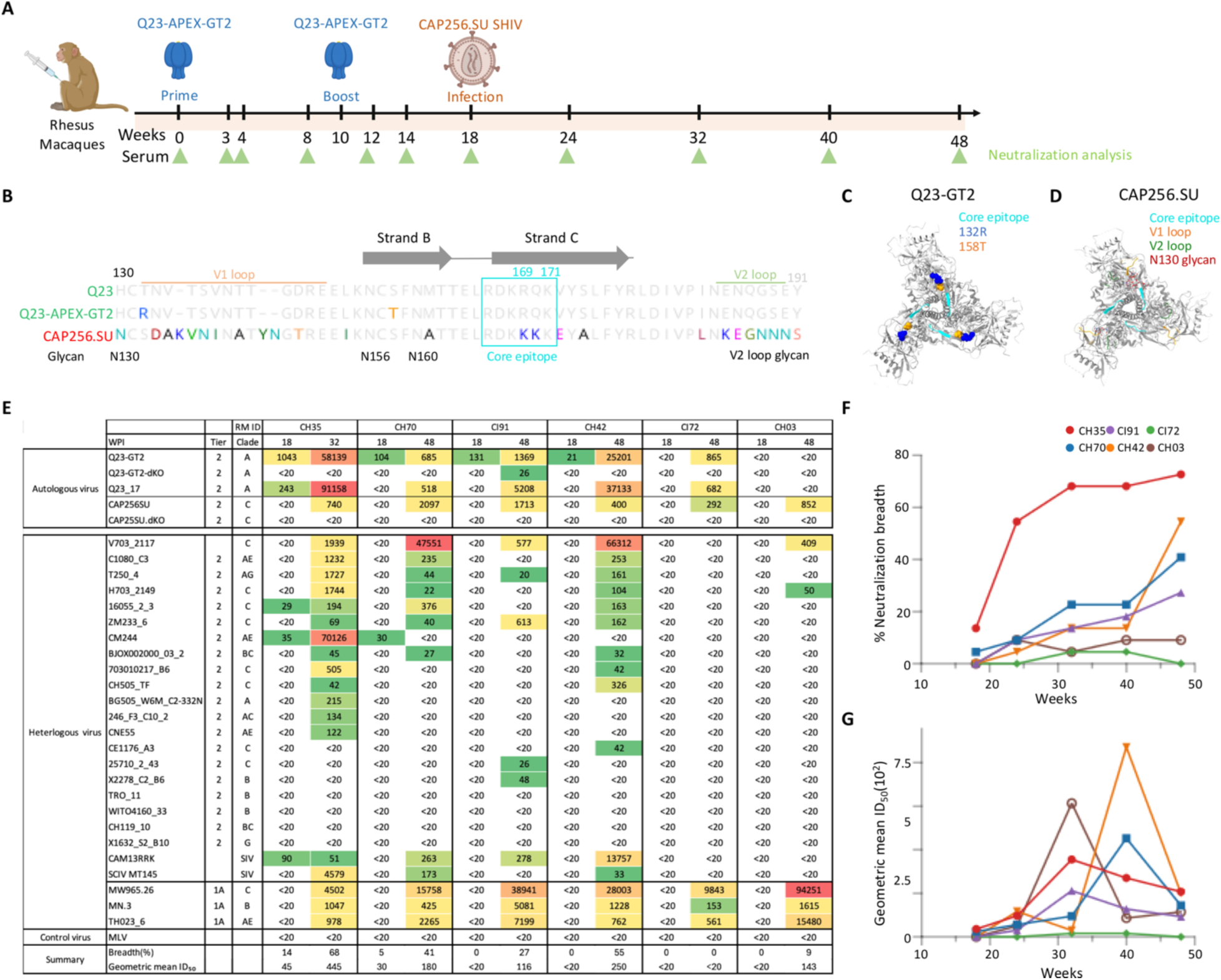
Serum neutralizing antibody responses following CAP256.SU SHIV infection in Q23-APEX-GT2–primed rhesus macaques. (**A**) Schematic of the immunization, infection, and sample collection timeline in rhesus macaques. The animals received the Q23-APEX-GT2 prime at week 0, a boost at week 10, and were infected with CAP256.SU SHIV at week 18. Serum samples were collected at weeks 0, 3, 4, 8, 12, 14, 18, 24, 32, 40 and 48 for neutralization assays. (**B**) Sequence alignment of the V1V2 regions from Q23, Q23-APEX-GT2, and CAP256.SU Env. V1 and V2 loops are indicated by lines, the V2-apex core epitope is highlighted by a box, and glycan sites are labeled. (**C** and **D**) Structural schematics of Env trimers: Q23-APEX-GT2 Env highlighting the core V2-apex epitope and germline-targeting mutations (C); CAP256.SU Env highlighting the V1V2 loop, core V2-apex epitope, and N130 glycan (D). (**E**) Serum neutralization summary table showing ID_50_ titers at week 18 (pre-infection) and weeks 32/48 (post-infection) for all animals against autologous viruses and a panel of heterologous viruses. (**F**) Longitudinal neutralization breadth curves for all animals against the 22-virus heterologous panel shown in panel (E). (**G**) Longitudinal geometric mean ID₅₀ curves for all animals against the 22-virus heterologous panel shown in panel (E). Following CAP256.SU SHIV infection, animals CH35, CH70, CI91, and CH42 developed serum neutralization breadth to varying degrees, with the most pronounced breadth observed in CH35.

We assessed longitudinal serum neutralization before and after CAP256.SU SHIV infection against autologous Q23-APEX-GT2 and CAP256.SU viruses and a heterologous panel of 22 viruses. Five of the six animals developed potent autologous neutralizing activity against the Q23-APEX-GT2 immunogen-matched virus, whereas all six exhibited potent serum neutralization against the CAP256.SU SHIV infection virus. Epitope-mapping analyses further showed that serum neutralization was directed almost exclusively to the V2-apex bnAb site in all six animals, indicating that the focused V2-apex specificity established by Q23-APEX-GT2 vaccination was boosted and retained following CAP256.SU SHIV infection (Fig. 1E and fig. S1).

Four of six animals (CH35, CH70, CI90, and CH42) subsequently developed broad heterologous tier 2 serum neutralization, neutralizing 27–68% of the 22-virus panel. The kinetics of breadth development varied among animals. CH35 developed broad neutralization rapidly, reaching peak neutralizing activity by week 32, just 14 weeks after CAP256.SU SHIV infection. In contrast, CH70, CI90, and CH42 reached peak breadth around week 48, 30 weeks after infection. All four animals developed potent serum neutralizing activity, with geometric mean ID₅₀ titers ranging from 116 to 445; CH35 exhibited the strongest response, with a geometric mean ID₅₀ of 445 (Fig. 1, E to G, and fig. S1).

The frequency and kinetics of breadth development were notable compared with previous studies of CAP256.SU or Q23.17 SHIV infection alone, in which broad V2-apex-directed serum neutralization developed less frequently and more slowly (*42, 61*). These findings support the premise that Q23-APEX-GT2 immunization established V2-apex-targeted B-cell populations that could be recalled following CAP256.SU SHIV infection, facilitating the development of potent serum neutralization and extending activity beyond the immunogen-matched virus. The differences in the magnitude and kinetics of breadth among animals may reflect variation in the Q23-APEX-GT2 vaccine-established B-cell repertoires and the capacity or compatibility of individual vaccine-elicited lineages to recognize and mature in response to CAP256.SU Env.

We next investigated whether the development of neutralization breadth was accompanied by viral evolution consistent with V2-apex bnAb-mediated immune selection pressure. Longitudinal sequencing of CAP256.SU viral populations revealed substitutions at V2 strand-C positions, including K169 and K171, as well as changes involving N160, R166, and other residues, across the four animals that developed serum neutralization breadth (fig. S2B). The extent and timing of these changes varied among animals. In CH35, a mutation at position K169 emerged by week 14 after infection with the K169E substitution becoming dominant by week 48, accompanied by an additional substitution at K171 (fig. S2B). Notably, cryo-EM analysis of the vaccine-elicited antibody CH35-Apex1.08, isolated before SHIV infection and in complex with Q23-APEX-GT2, identified K169 and K171 as antibody-contact residues (*32*). The emergence of substitutions at these positions is consistent with V2-apex-directed selection and suggests that antibody-mediated viral escape occurred during the development of neutralization breadth. Similar strand-C changes were observed in CH42, while CH70 and CI91 also developed substitutions at positions 169 and 171 (fig. S2B). In contrast, CH03 and CI72, which did not develop broad serum neutralization, lacked these changes, consistent with weaker or absent V2-apex-directed selection. Serum viral load measurements showed no sustained suppression across the cohort, indicating that the observed antibody responses and viral escape did not result in control of viremia (fig. S2A).

Collectively, these findings demonstrate that Q23-APEX-GT2 immunization can establish V2-apex bnAb precursor populations that are recalled following CAP256.SU SHIV infection and develop into responses associated with broad heterologous tier 2 serum neutralization. Four of six vaccinated animals developed serum breadth, with responses emerging as early as 14 weeks after infection. Thus, Q23-APEX-GT2 immunization priming establishes a foundation for the subsequent development of V2-apex serum bnAb responses following CAP256.SU SHIV infection. We next dissect the molecular features of these responses to determine how early B-cell recruitment, clonal expansion, and lineage development following Q23-APEX-GT2 priming shape the eventual acquisition of serum neutralization breadth.

### Q23-APEX-GT2–induced V2-apex autologous neutralization predicts broad serum neutralization after CAP256.SU SHIV infection

To determine whether Q23-APEX-GT2 elicited early B cell and antibody responses at priming influenced the development of serum neutralization breadth following CAP256.SU SHIV infection, we longitudinally analyzed serum polyclonal antibodies and B cell responses in lymph nodes and peripheral blood mononuclear cells (PBMCs) from all six Q23-APEX-GT2–immunized macaques (Fig. 2A). We first assessed serum samples collected through week 18, prior to SHIV infection, for autologous neutralization of Q23-APEX-GT2 virus. By week 12, five of six animals had developed measurable autologous serum neutralization (Fig. 2B and table S1). This neutralizing activity was directed against the V2-apex bnAb epitope, as neutralization was lost against a strand C double-knockout variant (R169E–K171E) that disrupts recognition by most V2-apex bnAbs (*29*) (Fig. 2B and table S1). The sera also exhibited enhanced neutralization of the N160K Q23-APEX-GT2 variant, which lacks the N160 glycan, a key component of the V2-apex bnAb epitope, while retaining dependence on strand C (table S1). These findings confirm focused priming of V2-apex– directed neutralizing responses, consistent with our previous report (*32*), although response magnitude varied among animals.

**Fig. 2.**
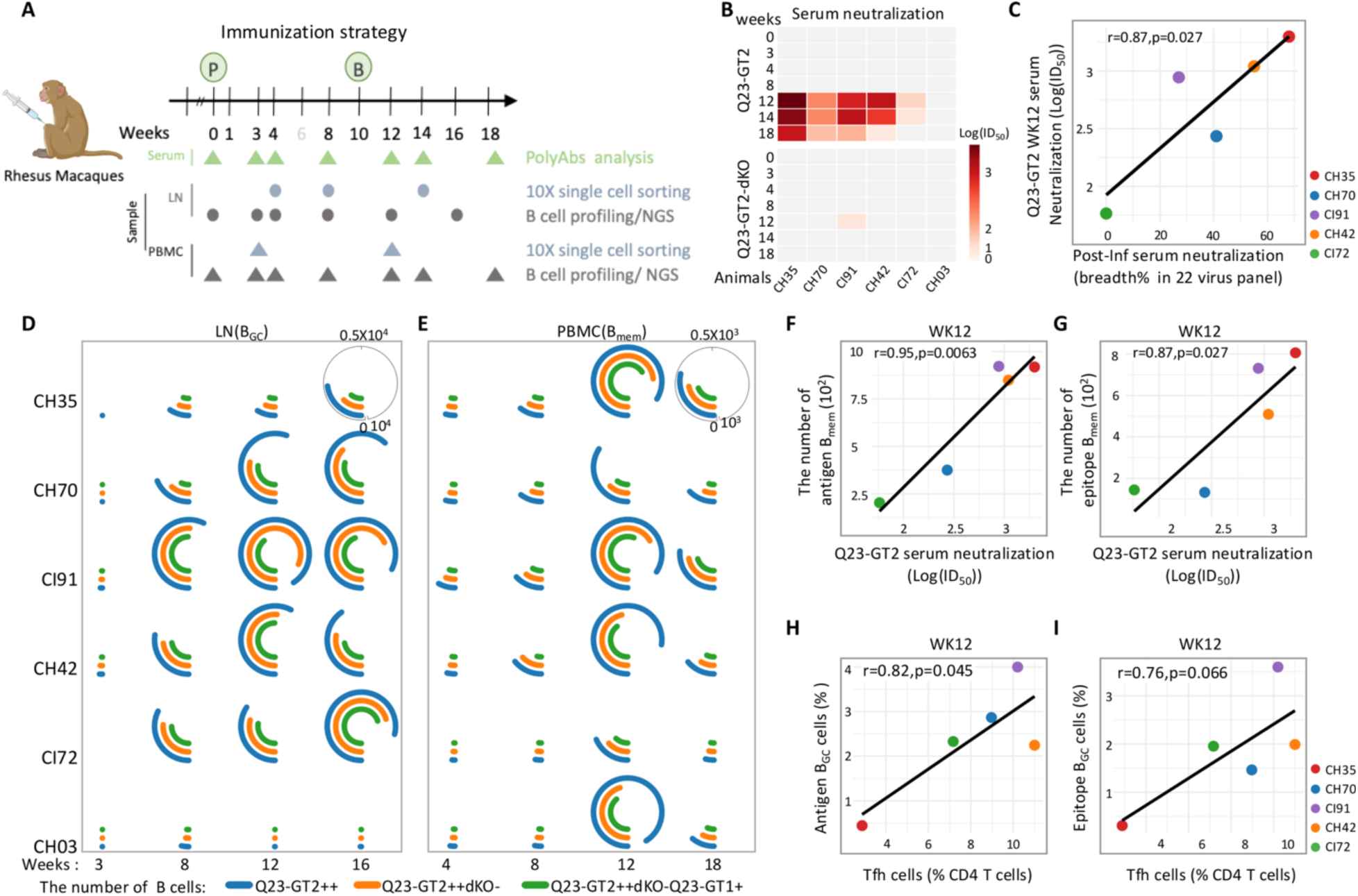
Longitudinal serum neutralizing antibody and B cell responses following Q23-APEX-GT2 immunization in rhesus macaques. (**A**) Immunization scheme and sample collection timeline. Rhesus macaques were primed at week 0 with 100 μg Q23-APEX-GT2 trimer formulated with SMNP adjuvant, administered subcutaneously across four injection sites (25 μg per site), following a two-week escalating-dose (DE) regimen. Animals were subsequently boosted at week 10 by bolus administration of 100 μg homologous trimer protein formulated with SMNP. Longitudinal sampling of serum, lymph nodes (LN), and peripheral blood mononuclear cells (PBMCs) was performed in parallel with functional analyses. (**B**) Longitudinal serum neutralization against homologous viruses. Heatmap summarizing ID₅₀ neutralization titers at weeks 0, 3, 4, 8, 12, 14, and 18 against pseudoviruses expressing Q23-APEX-GT2 (132R–158T) and its strand C dKO variant. Neutralization was abolished against dKO variants in Q23-APEX-GT2 backbones, confirming strand C dependency. (**C**) Correlation between week 12 Q23-APEX-GT2 specific autologous serum neutralization titers (Log_10_(ID₅₀)) and post-infection serum neutralization breadth against a heterologous 22-virus panel. (**D** and **E**) Semi-circular bar plots summarizing B cell subsets: GC B cell subsets in LNs (D) and memory B cell subsets in PBMCs (E). Subsets include antigen-specific (Q23-APEX-GT2⁺⁺), epitope-specific (Q23-APEX-GT2⁺⁺dKO⁻), and cross-reactive (Q23-APEX-GT2⁺⁺dKO⁻Q23-APEX-GT1⁺) populations. Colored arcs represent: blue, antigen-specific; orange, epitope-specific; green, cross-reactive. Arc length reflects the number of cells in each subset. (**F** and **G**) Correlation analyses between week12 serum neutralization titers (Log_10_ (ID_50_)) and the number of antigen specific (F) and epitope specific (G) memory B cells (B_mem_) in PBMCs at week12. (**H** and **I**) Correlation analyses between the frequency of Tfh cells and the frequencies of antigen-specific (H) and epitope-specific (I) germinal center B cells in lymph nodes at week12. Q23-APEX-GT2 immunization elicited robust antigen- and epitope-specific B cell responses in all animals, with PBMC memory B cell responses correlating with serum antibody activity.

Across the cohort, week 12 autologous serum neutralization titers were positively correlated with the development of serum neutralization breadth following SHIV infection (Fig. 2C; fig. S4, A and B). Of the five animals with detectable Q23-APEX-GT2 vaccine-induced autologous neutralization (CH35, CH70, CI91, CH42, and CI72), four (CH35, CH70, CI91, and CH42) subsequently developed serum neutralization breadth after SHIV infection. These four animals maintained detectable autologous neutralization through week 18, suggesting greater persistence of their vaccine-induced antibody responses. In contrast, CI72 exhibited weaker neutralization at week 12, which was no longer detectable by week 18, and did not develop comparable neutralization breadth following infection.

CH35 animal exhibited the strongest early response, with a week 12 autologous serum neutralization titer (ID₅₀ = 1,994) approximately fourfold higher than the cohort geometric mean (ID₅₀ = 497). Notably, CH35 also displayed the most pronounced evidence of cross-neutralization before infection. Its serum neutralized wild-type Q23.17, an activity not observed in the other animals except for transient neutralization in CH70 at week 12 and exhibited cross-neutralization of the heterologous CAM13RRK and CM244 viruses (table S1). This early serum cross-neutralization distinguished CH35 from the other animals and suggested more effective priming of antibodies capable of recognizing diverse V2-apex epitopes. Consistent with this possibility, CH35 rapidly developed the broadest and most potent serum neutralization response following CAP256.SU SHIV infection.

Together, these findings suggest that the magnitude, functional quality, and persistence of vaccine-induced early antibody responses may predict the subsequent development of serum neutralization breadth. We hypothesize that stronger early antibody responses reflect more effective recruitment and expansion of V2-apex–specific B cell lineages, some of which may persist in germinal centers and undergo further affinity maturation, while others differentiate into memory B cells and antibody-secreting plasma cells, potentially including long-lived plasma cells. Such responses could establish a durable pool of primed B cell lineages and antibodies that supports the development of broader neutralization following SHIV infection. Although the present data do not establish a causal relationship, they provide a rationale for investigating how early B cell recruitment, clonal selection, and persistence shape subsequent antibody maturation.

Overall, Q23-APEX-GT2 elicited V2-apex–focused autologous serum neutralizing responses in outbred macaques, with substantial inter-animal variation in response magnitude and breadth. This variation across the cohort provides an opportunity to dissect the B cell mechanisms that distinguish effective priming from responses that fail to progress toward broad serum neutralization.

### Antigen- and epitope-specific B cell responses

To further understand the immunological basis of differential neutralizing antibody responses following Q23-APEX-GT2 vaccination, we longitudinally quantified antigen-specific (Q23-APEX-GT2⁺ dKO⁺) and V2-apex–specific (Q23-APEX-GT2⁺ dKO⁻) germinal center (GC) B cells (B_GC_), memory B cells (B_mem_), antibody-secreting cells (ASCs), and GC T follicular helper (Tfh) cells in lymph nodes (LNs) and peripheral blood through week 18 (Fig. 2, D to I, and fig. S3) (*62*). We then assessed associations between these responses and autologous serum neutralization.

All animals developed antigen-specific GC B cell (CD38⁻CD71⁺) responses after priming, peaking at week 8 (0.2–2.5% of total B cells) and increasing further after the week 10 homologous boost (fig. S3, D and F). V2-apex–specific B_GC_ cells comprised 0.1–2.2% of the population. CI91 exhibited the highest antigen-specific B_GC_ frequencies and the greatest proportion of V2-apex– specific B_GC_ cells. CH35 had comparatively lower B_GC_ frequencies (fig. S3G). These differences likely reflect both biological variability and sampling constraints, as only 2–3 LNs were biopsied per animal at each time point. LN ASCs (CD27⁺CD38⁺) increased up to threefold, generally peaking at week 12, while GC-Tfh cells (PD-1^hiCXCR5⁺) expanded up to tenfold after priming, peaking at week 8 and stabilizing or increasing modestly after boosting (fig. S3, H and I).

Antigen- and epitope-specific B_mem_ frequencies in blood peaked at week 12 (0.1–1.3%) before contracting (fig. S3E). Notably, CH35 and CH03 developed the strongest circulating antigen-specific B_mem_ responses despite relatively low LN B_GC_ frequencies. Across animals, higher LN B_GC_ frequencies generally coincided with lower circulating B_mem_ frequencies, suggesting differences in the balance between ongoing GC responses and memory B cell differentiation (fig. S3F). Animals with stronger B_mem_ responses also tended to have marginally higher circulating ASC frequencies, consistent with coordinated memory and antibody-secreting cell responses (fig. S3, J and K).

To assess the quality of responses, we quantified antigen- and epitope-specific B cells per million lymphocytes and evaluated binding to the WT-derived Q23-SCT27 trimer (*31, 63, 64*). Animals with stronger LN antigen specific B_GC_ responses generally had more Q23-SCT27–binding, V2-apex–specific B_GC_ cells. In contrast, all animals developed substantial populations of circulating Q23-SCT27–binding B_mem_ cells. CH35 showed the strongest enrichment: at week 12, up to 78% of its V2-apex–specific B_mem_ cells recognized Q23-SCT27, compared with 29–66% in the other animals, and this enrichment persisted through week 18. This response coincided with unique ability of CH35 to neutralize WT Q23.17 (Fig. 2, D and E, and fig. S3, D and E). Thus, although most animals generated V2-apex–specific B cells capable of recognizing WT Q23, binding alone was insufficient for detectable WT neutralization, suggesting that additional antibody maturation or structural features were required for effective cross-neutralization.

The number of circulating antigen-specific and V2-apex–specific B_mem_ positively correlated with autologous serum neutralization titers against Q23-APEX-GT2, in animals where neutralization was detected, with significant associations at week 12 (Fig. 2, F and G, and fig. S4C). The correlations were generally stronger for total antigen-specific B_mem_ than for V2-apex–specific subsets, potentially reflecting contributions from responses targeting non–V2-apex epitopes. In contrast, LN B_GC_ frequencies did not significantly correlate with serum neutralization (fig. S4C). LN Tfh frequencies positively correlated with total and antigen-specific and V2-apex–specific B_GC_ frequencies, consistent with coordinated GC responses, but were not associated with LN or circulating ASC or B_mem_ frequencies (Fig. 2, H and I, and fig. S4, D and E).

Collectively, Q23-APEX-GT2 elicited robust antigen- and V2-apex–specific B cell responses across all six macaques. Circulating B_mem_ responses, in particular, correlated with autologous serum neutralization, which was itself associated with subsequent neutralization breadth following SHIV infection. These findings link early vaccine-induced B cell responses to serum neutralizing activity and support further investigation of how the magnitude, specificity, and maturation of primed B cell populations influence the development of broader neutralization.

### Early V2-apex B cell priming by Q23-APEX-GT2 predicts post-infection serum neutralization breadth

To understand the basis of serum V2-apex–directed neutralizing antibody responses, we asked whether these responses arise predominantly from the selective expansion of only a few long CDRH3 V2-apex bnAb lineages or from the activation of multiple distinct precursor lineages. Both mechanisms may contribute, with preferential expansion of selected clones shaping the magnitude and breadth of serum neutralization. To address this, we longitudinally isolated epitope-specific B cells from lymph node and PBMC compartments in all six Q23-APEX-GT2– immunized rhesus macaques to determine the contributions from B_GC_ and B_mem_ cells (Fig. 3A). V2-apex–specific IgG⁺ B cells were isolated by FACS from LN biopsies (axillary and inguinal) at weeks 4, 8, and 14 and from PBMCs at weeks 3 and 12, followed by single-cell and 10x Genomics-based sequencing of paired heavy and light chain repertoires to define clonal expansion, somatic hypermutation (SHM), and lineage relationships. These LN and PBMC time points were selected to maximize capture of major long-CDRH3 V2-apex lineages following the prime (week 0) and boost (week 10) in both germinal center and memory compartments (Fig. 3A and figs. S3A and S5A).

**Fig. 3.**
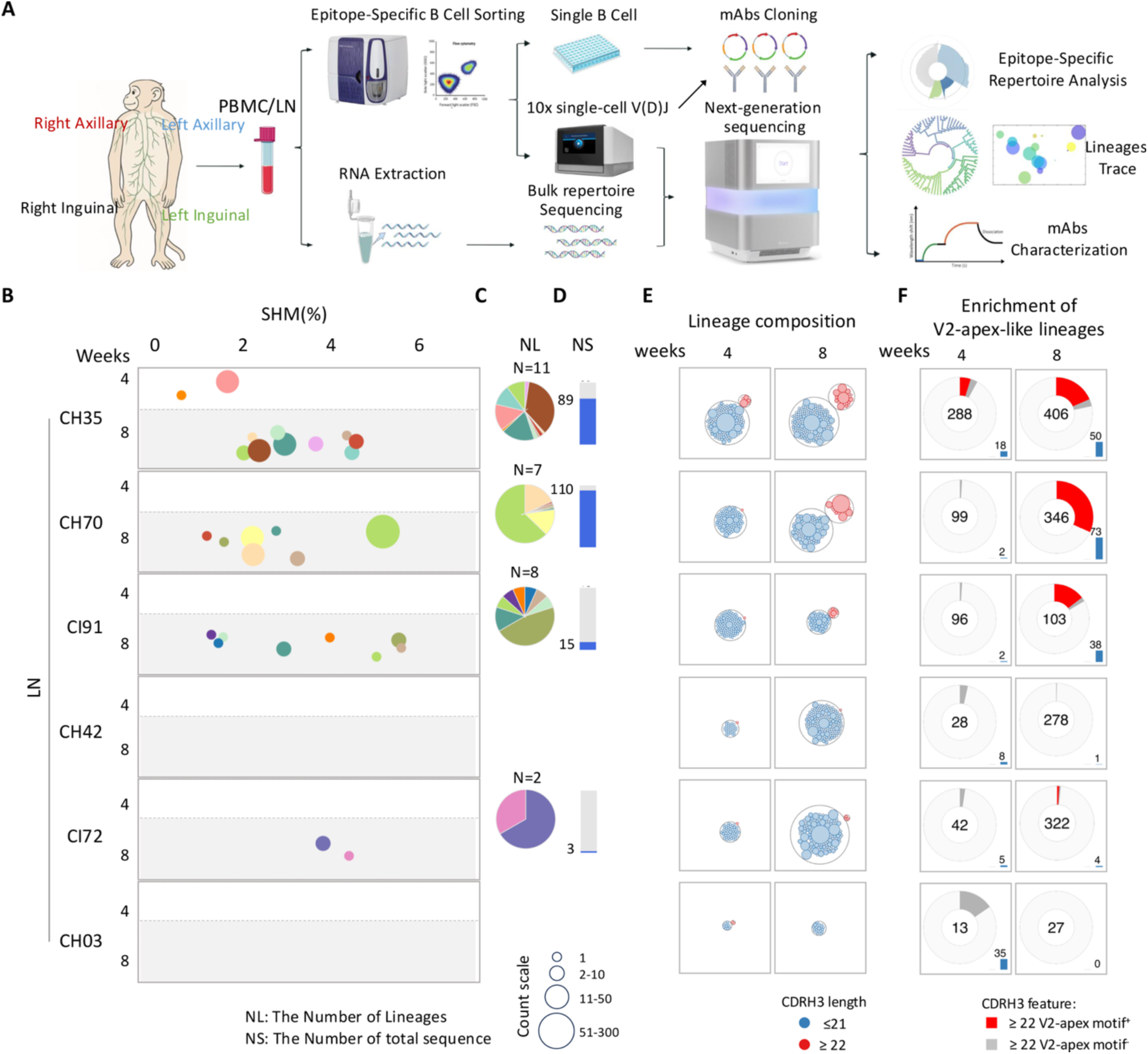
Early priming of epitope specific V2-apex long CDRH3 loop B cell lineages following Q23-APEX-GT2 immunization. (**A**) Schematic overview of experimental workflow. Lymph node (LN) biopsies (right/left axillary and inguinal) and peripheral blood mononuclear cells (PBMCs) were collected at multiple time points (pre-bleed, weeks 4, 8, 14, and 16/18). Epitope-specific IgG⁺ B cells were isolated via fluorescence-activated cell sorting (FACS), followed by single-cell mRNA extraction and Sanger sequencing. Bulk antibody repertoires from LN and PBMC samples, as well as 10x Genomics libraries from sorted epitope-specific IgG⁺ B cells, were generated for next-generation sequencing (NGS). Clonal lineage and repertoire analyses were performed to assess vaccine-induced B cell responses. Monoclonal antibodies (mAbs) were characterized via binding assays (BLI) and neutralization testing. (**B**) Bubble plots of epitope-specific V2-apex long CDRH3 loop B cell lineages identified from post-priming LN samples. Lineages were defined by CDRH3 lengths ≥22 amino acids together with conserved V2-apex Ab-associated motifs. The x-axis shows somatic hypermutation (SHM) levels; each row represents one animal. For each animal, the upper section shows early time points (week 4 LN), and the lower section shows later time points (week 8 LN). Each bubble represents a unique B cell lineage; bubble size reflects the number of clonotype members. Colors denote distinct lineages. (**C**) Pie charts summarizing the distribution of epitope-specific V2-apex long CDRH3 loop B cell lineages in post-priming LN samples. Each sector represents one lineage, and the area of each sector is proportional to the number of sequences within that lineage. N denotes the number of unique lineages identified. (**D**) Bar plots showing the total number of V2-apex Ab-like clonotypes per animal in LN. (**E** and **F**) Lineage composition based on CDRH3 length and motif usage from epitope-sorted IgG⁺ B cells analyzed via 10X Genomics single-cell sequencing. (E) Circle packing plots show lineage distribution by CDRH3 length. Red circles indicate CDRH3 ≥22 amino acids; blue circles indicate CDRH3 <22 amino acids. Filled circles represent expanded lineages; bubble size reflects number of clonotype members. (F) Donut charts show the proportion of lineages with CDRH3 ≥22 amino acids that contain conserved V2-apex bnAb motifs (YYD, DY, DDY, (E|D)(E|D)Y, and (E|D)DDY; red) versus those without motifs (gray). Numbers within donuts indicate total sequences. Bar plots show fold change of CDRH3 ≥22 amino acid lineages in immunized rhesus macaques relative to naïve animals. Q23-APEX-GT2 successfully recruits multiple V2-apex long CDRH3 loop B cell precursor lineages, but the magnitude, diversity, and early expansion of these lineages vary across animals.

Distinct B cell clonal lineages bearing long CDRH3 loops (≥22 amino acids) and genetic hallmarks of V2-apex bnAbs—including characteristic D-gene usage and conserved anionic CDRH3 motifs (e.g., YYD, DY, DDY, (E/D)(E/D)Y, and (E/D)DDY)—were detected in the LNs of all six Q23-APEX-GT2-immunized animals by week 14, consistent with effective priming following two immunizations (fig. S7A). These lineages were enriched for long CDRH3 loops and preferentially used IGHD3-15 (fig. S5B), consistent with our previous isolation of V2-apex bnAb-like monoclonal antibodies from week 14 LNs (*32*). The lineages accumulated up to ∼6% heavy-chain somatic hypermutation (SHM), with levels varying across animals.

Long-CDRH3 V2-apex bnAb-like lineages were detected as early as week 4 in only one animal (CH35) and by week 8 in four of six animals (CH35, CH70, CI91, and CI72), with 2–11 distinct lineages identified per animal (Fig. 3B and fig. S6A). Notably, three of these four animals (CH35, CH70, and CI91) subsequently developed broad serum neutralization following SHIV infection. Across the weeks 4 and 8 LN sampling time points, these three animals exhibited greater V2-apex lineage diversity (11, 7, and 8 distinct lineages, respectively) and higher numbers of recovered epitope-specific long-CDRH3 B cells (89, 110, and 15, respectively) than CI72 (Fig. 3, B to D, and fig. S6A), which had only two distinct lineages and three recovered B cells and did not develop serum bnAbs after infection. These findings suggest that the magnitude and diversity of early V2-apex B cell priming may influence subsequent bnAb development. CH42 represented a notable exception: despite developing serum bnAb breadth following SHIV infection, no long-CDRH3 V2-apex lineages were detected in its week 4 or week 8 LN samples. This absence may reflect limited sampling rather than a failure of priming, as only 2–3 LNs (out of ∼500 LNs in a monkey) were biopsied per animal at each time point. Furthermore, CH42 developed high early autologous neutralizing antibody titers, another correlate of subsequent serum bnAb development, potentially reflecting productive responses in unsampled lymphoid tissues. The subsequent isolation of V2-apex bnAb lineages from CH42 (week 14 LN) following homologous boosting further supports the possibility that these lineages were likely present but undetected during early LN sampling (fig. S6A). Together, these findings suggest that efficient early priming and expansion of diverse V2-apex bnAb-like B cell lineages by Q23-APEX-GT2 may establish an immunological trajectory favorable for the development of serum bnAb activity following SHIV infection.

Following homologous Q23-APEX-GT2 boosting, long-CDRH3 V2-apex bnAb-like lineages were detected in week 14 LN samples (4 weeks post-boost) from all six animals, with 8, 12, 3, 2, 2, and 3 lineages identified in CH35, CH70, CI91, CH42, CI72, and CH03, respectively. Consistent with the priming phase, CH35, CH70, and CI91 exhibited more pronounced elicitation of diverse prototype V2-apex bnAb lineages and greater clonal expansion, with 47–200 long CDRH3 V2-apex B cells recovered per animal. Across priming and homologous boosting, these three animals, which subsequently developed serum neutralization breadth, each exhibited 10–19 distinct long-CDRH3 V2-apex lineages in LNs (fig. S6, A and B). Overall, homologous boosting elicited detectable V2-apex bnAb-like lineages in all six animals, but the magnitude and diversity of these responses varied and were associated with subsequent neutralization outcomes.

A similar trend was observed in PBMCs, where long-CDRH3 V2-apex memory B cell clones were first detected at week 3 in only two animals (CH35 and CH03) and expanded in diversity and SHM by week 12 (fig. S6A). Across animals, 1-7 unique long-CDRH3 V2-apex lineages were identified in the PBMC memory compartment, with CH35 again showing the greatest diversity (n = 7) (fig. S6, A and B). As expected, the size of expanded long-CDRH3 clones were substantially less in PBMCs than in LNs, consistent with ongoing germinal center activity expansion in lymphoid tissue. Notably, in 4 out of 6 animals, we detected at least one long-CDRH3 lineages that were shared between LN and PBMC compartments within individual animals, indicating coordinated clonal expansion and memory seeding (fig. S6, A and B).

To define the composition of epitope-specific B cell repertoires, we analyzed lineage distributions according to CDRH3 length and motif usage. In lymph nodes, long-CDRH3 lineages (≥22 amino acids) were readily elicited, with several undergoing substantial clonal expansion (Fig. 3, E and F, and fig. S6D). Relatively shorter CDRH3 lineages (≤21 amino acids) were also detected, with some showing substantial expansion. Notably, CH35, CH70, and CI91 exhibited higher proportions of long-CDRH3 lineages encoding conserved V2-apex bnAb-associated motifs before boosting, and these responses were maintained after boosting. In contrast, CH03 showed a marked increase in the proportion of these lineages only after boosting (Fig. 3F and fig. S6D). In PBMCs, shorter CDRH3 lineages predominated and were generally distributed across numerous, less-expanded clones. Nevertheless, CH35 and CH03 remained distinguished by higher frequencies of long-CDRH3 lineages bearing V2-apex bnAb-associated motifs (fig. S6E).

Overall, Q23-APEX-GT2 consistently primed V2-apex bnAb precursors across animals, but the magnitude and diversity of these responses varied. Animals that developed serum neutralization breadth (CH35, CH70, and CI91) exhibited greater lineage diversity and clonal expansion during priming, suggesting that robust early V2-apex B cell responses may establish a favorable trajectory for bnAb development following SHIV infection.

### Dominant expanded long-CDRH3 V2-apex lineages drive serum neutralization

To determine the full magnitude of long-CDRH3 V2-apex bnAb precursor priming, track their longitudinal expansion and maturation, and evaluate their contribution to serum neutralization, we surveyed epitope-specific V2-apex B cell lineages identified through epitope-specific sorting in longitudinal lymph node and PBMC samples using bulk next-generation sequencing (NGS) (Fig. 4, A to D and fig. S7). Analyses were performed across six LN time points (weeks 0, 3, 4, 8, 12 and 16) and seven PBMC time points (weeks 0, 3, 4, 8, 12, 14 and 18). Week 0 samples established baseline lineage frequencies, weeks 3-8 captured post-prime responses, and weeks 12-18 reflected post-boost dynamics. We reasoned that integrating dense longitudinal sampling with quantitative NGS across anatomical compartments would enable a more accurate assessment of lineage expansion, evolution, and functional contribution to serum neutralization. Indeed, quantitative lineage tracking provides a more comprehensive and less biased view of B cell dynamics than single-B cell isolation alone, which may under-sample key clones within specific compartments.

**Fig. 4.**
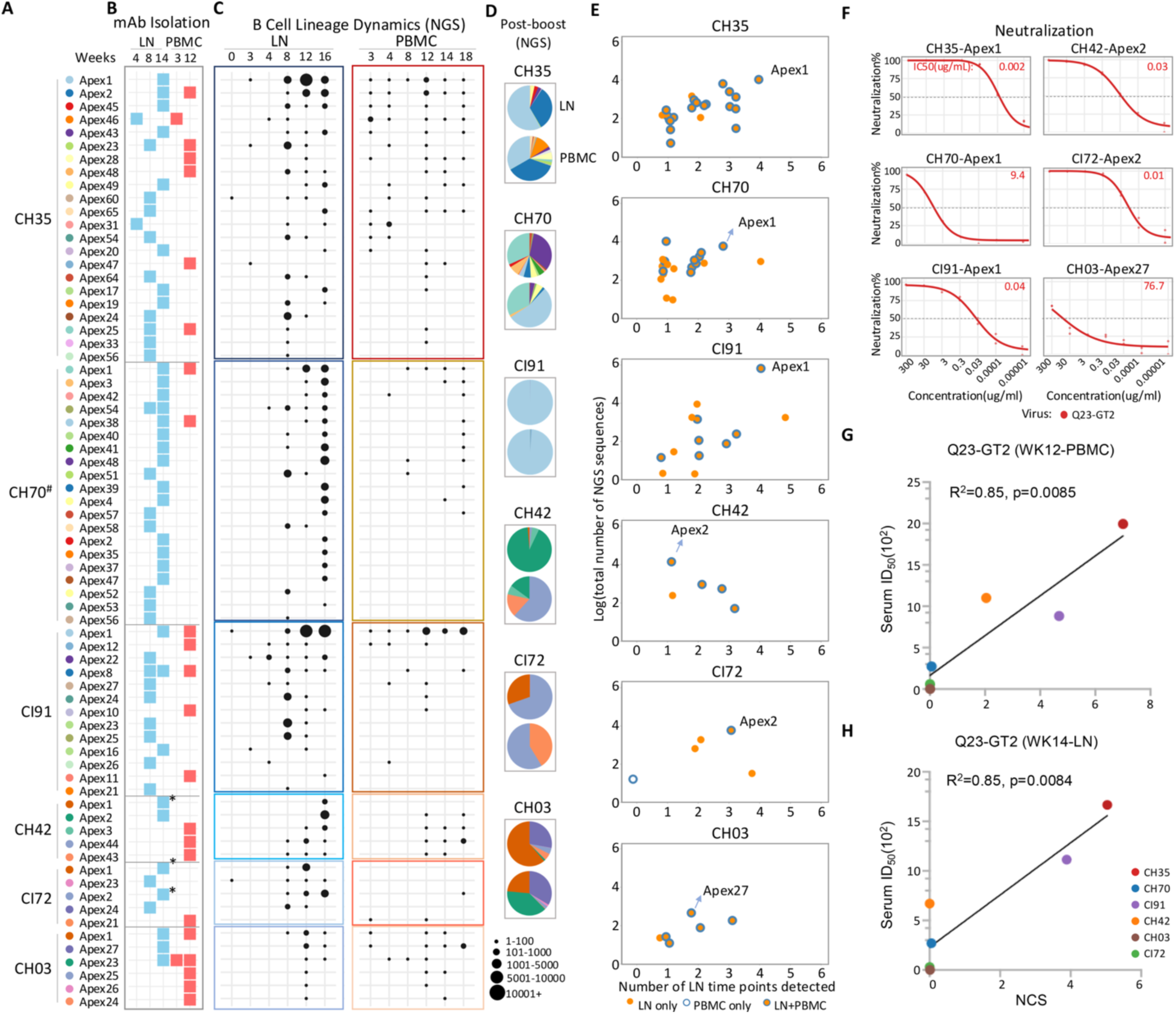
Longitudinal B cell clonal dynamics and neutralization characteristics of V2-apex long CDRH3 loop B cell lineages. (**A**) V2-apex long CDRH3 loop B cell lineages analyzed in this figure. All lineages were identified following Q23-APEX-GT2 priming and homologous boosting. #: For week 14 LN sample, CH70 is from week 16 LN; all others are from week 14. (**B**) Sampling timepoints at which monoclonal antibodies (mAbs) were isolated from each lineage. Blue indicates lymph node (LN) compartments (weeks 4 and 14); red indicates peripheral blood mononuclear cell (PBMC) compartments (weeks 3 and 12). (**C**) Bubble plot showing longitudinal dynamics of epitope-specific lineages identified in panel (A), based on bulk repertoire NGS. Lineages are grouped by animal and compartment (colored boxes). The x-axis shows timepoints: week 0, 3, 4, 8, 12, and 16 for LNs; week 0, 3, 4, 8, 12, 14, and 18 for PBMCs. Bubble size reflects the number of clonotypes within each lineage. (**D**) Pie charts illustrating lineage distribution in post-boost NGS data. Dominant lineages were observed in each animal: CH35-Apex1 (LN and PBMC), CH70-Apex1 (LN and PBMC), CI91-Apex1 (LN and PBMC), CH42-Apex2 (LN and PBMC), CI72-Apex2 (PBMC), CH03-Apex1 and Apex27 (LN and PBMC). (**E**) Dot plot showing expansion of all V2-apex long CDRH3 loop B cell lineages across animals. The x-axis indicates the number of LN time points at which each lineage was detected, and the y-axis represents the total number of sequences identified by bulk NGS. Orange-filled dots indicate lineages detected in LNs, blue-outlined dots indicate lineages detected in PBMCs, and orange-filled dots with blue outlines indicate lineages detected in both LN and PBMC compartments. Representative lineages are labeled. These representative lineages exhibited both substantial clonal expansion, sustained detection across multiple LN time points and distribution across both LN and PBMC, highlighting their dominance within the repertoire. (**F**) Neutralization titration curves of dominant lineage mAbs against autologous viruses (Q23-APEX-GT2). (**G** and **H**) Scatter plot showing the relationship between the Neutralization Contribution Score (NCS) and serum neutralization ID_50_ at week 12 PBMC(G) and WK14 LN(H). Serum ID₅₀ values and lineage abundances were derived from the corresponding sampling time point and compartment (week 12 PBMCs or week 14 LNs). NCS was calculated as 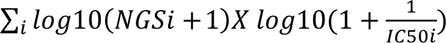, where NGS represents lineage abundance determined by bulk BCR sequencing and IC_50_ represents the neutralization potency of the corresponding monoclonal antibody. Solid line indicates linear regression. Following Q23-APEX-GT2 immunization, dominant V2-apexs Ab-like B cell lineages emerged in all animals with varying degrees of clonal expansion and neutralization potency. Integration of lineage abundance with monoclonal antibody activity demonstrated that preferential expansion of productive lineages was strongly associated with serum neutralization.

NGS revealed expansion of long-CDRH3 lineages in all animals, consistent with those identified by epitope-specific B cell sorting, indicating successful priming and subsequent trafficking across lymphoid tissues. These lineages were detectable as early as week 3 in LN samples and peaked post-prime by week 8, when numerous lineages were observed (Fig. 4C and fig. S7A). Following the homologous boost, one to two dominant lineages emerged in most animals by week 12, exhibiting substantial expansion consistent with recall responses, and most of the lineages remained detectable through week 16 (six weeks post-boost) (Fig. 4, C and D, and fig. S7). Most lineages were also detectable in the PBMC compartment, although their expansion was markedly lower than in LNs, suggesting that only a fraction of each lineage was seeded into the memory pool—consistent with the antigen-specific B cell data described above (Fig. 2D and fig. S7). While many lineages appeared early in PBMC (week 3) without corresponding detection in LN samples, this likely reflects sampling limitations, as LN biopsies represent individual nodes whereas PBMC-derived B cells capture contributions from the broader lymphatic network. Importantly, substantial overlap between LN and PBMC repertoires across animals supports shared lineage origins and active trafficking. Moreover, the size of dominant clonal lineages in LNs showed a positive association with memory representation in PBMCs (Fig.4, A to D and fig. S7).

Longitudinal analyses revealed that one or two dominant long-CDRH3 lineages typically drove the response in each animal and were shared across compartments (Fig. 4E and fig. S8). MAbs derived from dominant lineages in all five animals that developed Q23-APEX-GT2-specific serum neutralization by week 12 potently neutralized the autologous virus (Fig. 4F and fig. S9A). Together, these findings suggest that expanded long-CDRH3 lineages largely account for serum neutralization activity, with a limited number of dominant bnAb lineages driving the response.

To quantitatively assess the contribution of long CDRH3 lineages to serum neutralization, we developed a parameter-free Neutralization Contribution Score (NCS) integrating the neutralization potency (IC_50_) of lineage-derived IgGs with their expansion measured by bulk NGS (Fig. 4, G and H, and fig. S9, D and E). The NCS showed a strong positive correlation with serum neutralization ID_50_ titers at week12 (Pearson r=0.924, R^2^=0.85, P=0.0085), with a comparable association observed at week14. These findings support the notion that the magnitude of early lineage expansion, coupled with affinity maturation, is a key determinant of serum neutralization breadth and potency.

Together, these findings establish that vaccine-elicited serum neutralization is primarily driven by the preferential expansion and maturation of a small number of long-CDRH3 V2-apex bnAb lineages, highlighting the importance of early precursor recruitment and clonal dominance in shaping effective bnAb responses.

### Recall of V2-apex long CDRH3 B cell lineages in CH35 monkey upon CAP256.SU infection

To investigate the recall and maturation of Q23-APEX-GT2–primed long-CDRH3 V2-apex B cell lineages following CAP256.SU SHIV infection, we performed longitudinal bulk NGS analysis of lymph node and PBMC samples from macaque CH35. We selected CH35 because it exhibited the strongest priming response and subsequently developed the broadest serum neutralization following infection. Multiple long-CDRH3 lineages detected after vaccination were strongly recalled after infection, confirming effective re-engagement of vaccine-primed clones. Recall was evident by weeks 6-14 post-infection (weeks 24 and 32), with rapid expansion in LNs consistent with germinal center–driven maturation and subsequent dissemination into the circulating memory pool (Fig. 5A and fig. S10A). Most lineages peaked between weeks 24 and 32, although kinetics varied by lineage and compartment, indicating heterogeneous maturation trajectories (Fig. 5A and fig. S10A). While dominant clones were generally conserved across compartments, shifts in lineage frequencies revealed ongoing clonal competition and selection, with some previously minor lineages expanding substantially and reshaping the post-vaccination clonal landscape (Fig. 5, A and B and fig. S10A).

**Fig. 5.**
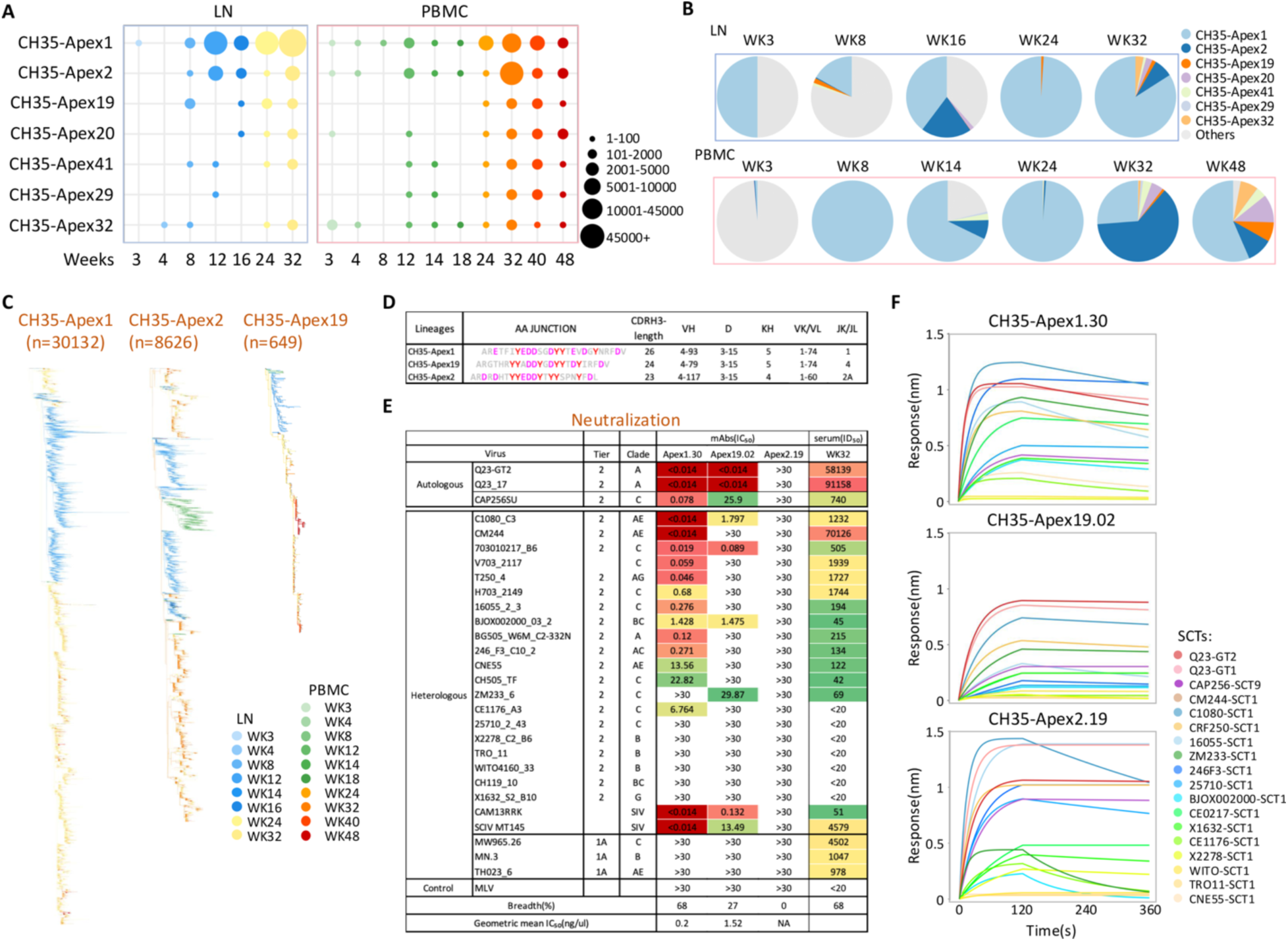
Lineage recall and maturation following CAP256.SU SHIV infection in CH35. (**A**) Temporal and spatial distribution of recalled lineages across lymph node (LN) and PBMC compartments. Each bubble represents a lineage at a specific time point, with size proportional to the number of sequences. CH35-Apex1 exhibited dramatic expansion post-infection. (**B**) Pie charts showing the percentage distribution of all isolated lineages in LN (post-prime: weeks 3 and 8; post-boost: week 16; post-infection: weeks 24 and 32) and PBMCs (post-prime: weeks 3 and 8; post-boost: week 14; post-infection: weeks 24, 32, and 48). Lineages recalled after CAP256.SU infection are color-coded. CH35-Apex1 dominated post-infection responses and underwent substantial clonal expansion. (**C**) Phylogenetic trees of representative V2-apex–specific lineages from CH35: CH35-Apex1, CH35-Apex2, and CH35-Apex19. *n* indicates the total number of clonotypes per lineage. Branch colors indicate sampling timepoints: blue/green for pre-infection, yellow/orange/red for post-infection. These lineages show clonal diversification and persistence across timepoints, with CH35-Apex1 demonstrating the most pronounced expansion and maturation from priming through post-infection stages. (**D**) Table summarizing heavy-chain CDRH3 amino acid junction sequences, CDRH3 length, V(D)J gene usage for heavy chains, and VJ gene usage for light chains across representative lineages. (**E**) Neutralization summary comparing the IC_50_ values of post-infection mAbs derived from the Apex1, Apex2, and Apex19 lineage against autologous and heterologous viruses, alongside CH35 week 32 serum ID_50_ titers. CH35-Apex1, which underwent extensive clonal expansion, exhibited the broadest neutralization breadth and closely mirrored serum neutralization potency, indicating it is a major contributor to serum neutralizing activity. (**F**) BLI binding curves showing Env SCT binding profiles for a subset of viruses from panel (E) for Apex1, 2, and 19. All lineages displayed broadly heterologous binding profiles. Multiple V2-apex–specific lineages were recalled following CAP256.SU SHIV infection. Among these, CH35-Apex1 showed the greatest clonal expansion and neutralization breadth, dominating post-infection responses.

Seven major long-CDRH3 lineages (CH35-Apex1, CH35-Apex2, CH35-Apex19, CH35-Apex20, CH35-Apex41, CH35-Apex29 and CH35-Apex32) were tracked to assess affinity maturation, expansion, and turnover. All accumulated increasing somatic hypermutation after infection, consistent with sustained GC activity, but the magnitude of maturation differed markedly (Fig. 5C and fig. S10, B to D). The CH35-Apex1 lineage exhibited sustained expansion and dominated both LN and PBMC compartments, whereas CH35-Apex2 showed pronounced expansion in blood at week 32 followed by contraction, suggesting likely lower fitness under antigenic selection (Fig. 5B and fig. S10, E and F). These findings indicate that SHIV infection not only recalls vaccine-primed precursors but also drives iterative mutation and selection that refine clonal hierarchy.

Monoclonal antibodies from each recalled lineage were evaluated longitudinally for epitope specificity, binding breadth, and neutralization against autologous and heterologous viruses. All seven retained V2-apex specificity (tables S2 and S3). Three neutralized autologous viruses, two displayed cross-neutralization limited to a few heterologous viruses, and one lineage, CH35-Apex1, evolved into a highly broad and potent bnAb that likely accounted for the majority of serum neutralizing activity (Fig. 5E, and table S3). In contrast, the strongly recalled CH35-Apex2 lineage expanded substantially and overtook CH35-Apex1 in blood by week 32 yet remained entirely non-neutralizing (Fig. 5E and table S3). Despite cross-reactive binding to diverse heterologous trimers comparable to CH35-Apex1, CH35-Apex2 failed to neutralize even the autologous virus (Fig. 5F and table S2). We isolated and characterized multiple CH35-Apex2 lineage members from week 32 post-infection time point and found that, despite extensive affinity maturation, they retained a non-neutralizing phenotype (table S3). The CH35-Apex2 lineage recall by SHIV Env implies recognition of a native-like trimer in an infectious virion, but the absence of functional neutralizing activity suggests unfavorable Env engagement incompatible with neutralization. CH35-Apex2 therefore represents a “born-wrong” V2-apex lineage—despite exhibiting canonical V2-bnAb–like precursor features (Fig. 5D), it expands and matures but is intrinsically incapable of acquiring neutralizing function.

Our longitudinal lineage tracking demonstrates that CAP256.SU infection efficiently recalled multiple epitope-specific lineages seeded by Q23-APEX-GT2 immunization. These lineages accumulated additional somatic hypermutation, and some exhibited enhanced neutralization following infection, demonstrating that Q23-APEX-GT2 germline-targeting priming can establish bona fide bnAb precursors capable of further maturation upon antigenic re-exposure. However, not all bnAb-like precursors progressed toward breadth; some underwent substantial clonal expansion yet remained non-neutralizing. These findings highlight a key advantage of recruiting a diverse precursor repertoire during priming: because individual lineages differ in their capacity to recognize and neutralize diverse viruses, engaging multiple candidate lineages may increase the likelihood of eliciting breadth. Thus, although germline-targeting strategies can seed diverse bnAb precursor lineages, their eventual success depends on selective clonal expansion and maturation along trajectories that confer broad neutralization.

### Structural basis of heterologous neutralization breadth

To investigate the basis for differences in heterologous neutralization breadth of lineages recalled and expanded by CAP256.SU SHIV infection, we determined the cryo-EM structures of antigen-binding fragments (Fabs) from CH35-Apex2.03 and CH35-Apex19.01 in complex with the Q23-APEX-GT2 trimer (fig. S11), and compared them to the complex structure of an earlier pre-infection lineage member of CH35-Apex1.30, CH35-Apex1.08 (*32*).

We first compared the binding orientation and angle of approach for CH35-Apex1.08, Apex2.03, and Apex19.01 Fabs by alignment of the Env trimer from each structure (Fig. 6A). While all three lineages recognized the V2-apex with highly similar orientations of their respective heavy and light chains, CH35-Apex2.03 exhibited an uncharacteristically low angle of approach (*61*) that was ∼16 degrees closer to the horizontal apex plane than CH35-Apex1 and Apex19.01. As a result, the CH35-Apex2.03 heavy chain made substantial interactions with the V3 region, including the N-terminal beta-strand (residues 305-308) and the loop immediately preceding the GDIR motif (Fig. 6B and fig. S12A). In contrast, CH35-Apex1.08 and Apex19.01 Fabs were nearly superimposed with each other (Fig. 6A), had angles of approach like human and rhesus V2 apex bnAbs with axe-like and combined modes of recognition (fig. S12B), and did not contact residues outside of V1V2 (Fig. 6B). Overall, this analysis suggests the substantial off-target V3 component of the CH35-Apex2.03 epitope contributed to its inability to neutralize autologous or heterologous HIV strains.

**Fig. 6.**
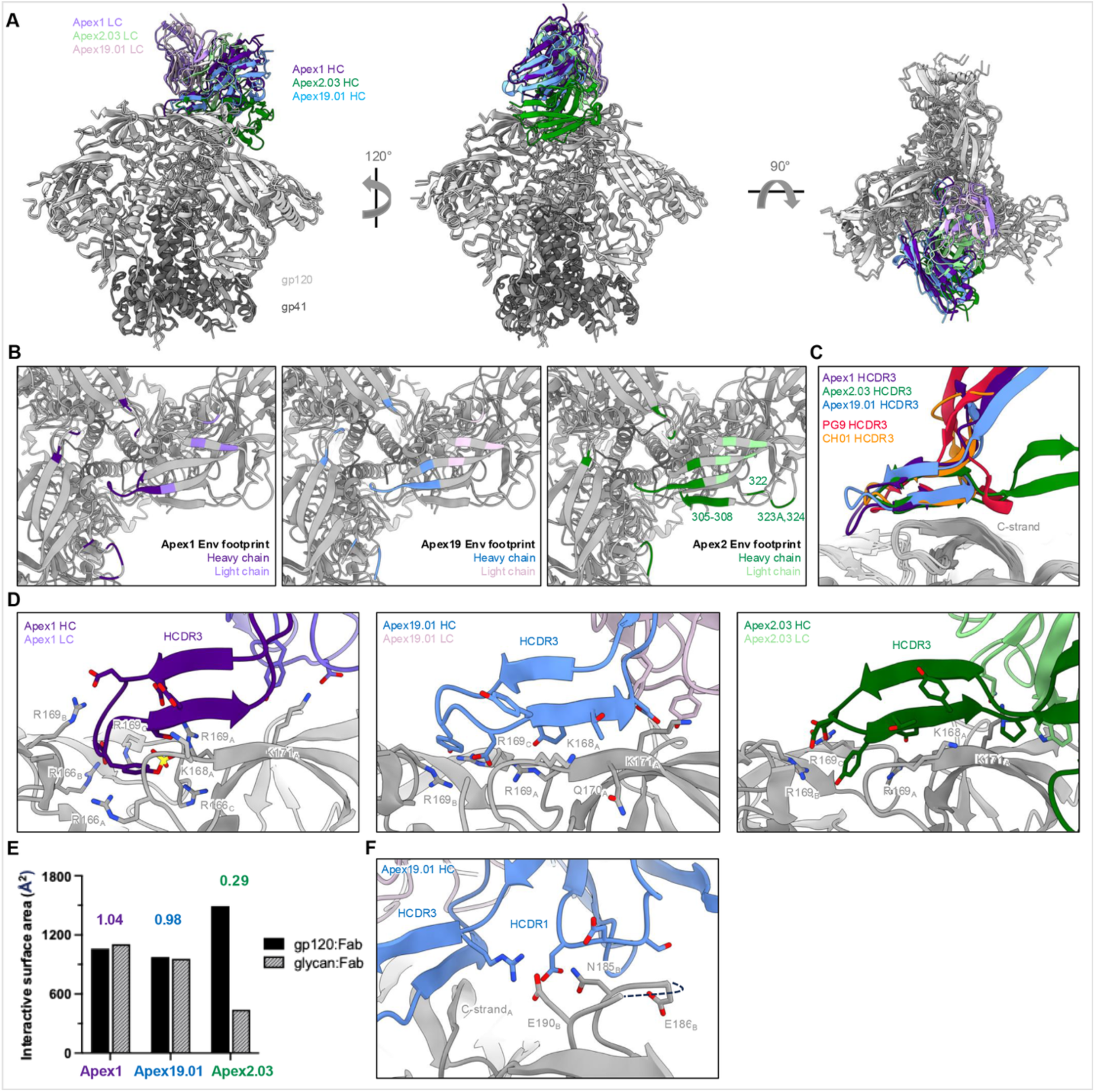
Structural basis of V2 apex recognition by SHIV-CAP256SU recalled lineages. (**A**) Orthogonal views for the superimposed cryo-EM structures of CH35-Apex1.08, Apex2.03, and Apex19.01 antigen-binding fragments (Fabs) in complex with Q23-APEX-GT2 Env trimer. Env gp120 and gp41 in each structure are colored light gray and tungsten, respectively. Fab heavy (HC) and light chains (LC) are colored according to lineage. (**B**) The 4-Å Env footprints of CH35-Apex1.08, Apex2.03, and Apex19.01 Fabs are mapped onto each respective trimer complex and colored by heavy or light chain contact. The remaining Env surface is shown in gray, and all other components of the structure are omitted for clarity. Top view of the trimer is presented. (**C**) HCDR3 interface side view of the structures of CH35-Apex1.08, Apex2.03, and Apex19.01 superimposed with the Env complex structures of human Fabs PG9 (PDB-7T77) and CH01 (PDB-9DHW). Other components of each Fab structure are omitted to highlight HCDR3 structure and positioning. (**D**) Further expanded HCDR3 interface view of CH35-Apex1, Apex2.03, and Apex19.01 to highlight interactions with V2 apex C-strand residues. Epitope:paratope residues are depicted in stick representation with carbon atoms colored by chain, nitrogen atoms colored dark blue, oxygen atoms colored bright red, and sulfur atoms colored yellow. V2 apex residue contacts are labeled. (**E**) Quantification of CH35-Apex1, Apex2.03, and Apex19.01 interactive surfaces are partitioned by gp120 protein and glycan interfaces. The ratio of glycan:gp120 surface area is listed above each pair of columns. (**F**) CH35-Apex19.01 interactions with hypervariable V2 loop from neighboring protomer_B_. Epitope:paratope residues are depicted similarly to panel (D). Three residues in the V2 loop (187, 188, 188A) that become disordered and cannot be accurately modeled are indicated with a dashed line.

Despite the two distinct approaches to the trimer apex, all three CH35 lineages had highly similar V1V2 contact residues, especially within the C-strand (Fig. 6B). This was achieved through insertion of elongated HCDR3s with convergent axe-like topology that mainchain bonded with the C-strand, like human bnAbs PG9 and CH01 (*51, 52, 61*) (Fig. 6C and fig. S12C). Consistent with its lower binding angle, the CH35-Apex2.03 HCDR3 was inserted horizontally to the trimer apex plane to reach the C-strand, thus revealing how a non-classical apex-binding angle is compatible with C-strand dependance (tables S2 and S3). CH35-Apex1.08, Apex2.03, and Apex19.01 each had conserved contacts with C-strand residues K168 and K171 from the primary recognized protomer_A_ and R169 from all protomers (Fig. 6D). Paratope chemistries were also largely conserved among the CH35 lineages to bind the Arg and Lys residues in the C-strand, which was primarily mediated by anionic residues forming salt bridges with the positively charged Arg/Lys amine groups and aromatic residues stabilizing the extended Arg/Lys aliphatic chains. CH35-Apex1.08 was unique in its recognition of R166 from all protomers, which was achieved in part by insertion of the tyrosine-sulfated HCDR3 loop tip into the trimer apex hole (Fig. 6D, left). Thus, each CH35 lineage had distinct patterns of heterologous neutralization despite all exhibiting canonical C-strand recognition, suggesting this to not be a significant contributing factor to their disparate neutralization breadths – although, it is likely that the additional binding energy mediated by the posttranslational tyrosine sulfation modification in the CH35-Apex1 lineage contributes to the higher geomean IC_50_ titer (Fig. 5E) (*65, 66*).

Another component of the V2 apex epitope is the apical glycan shield, which is most critically composed of *N*-linked glycans at Env residues 130, 156, and 160 (*29, 51–54, 64*). We have previously found that antibodies elicited by a single homologous prime and boost immunization with Q23-APEX-GT2 can neutralize heterologous viruses if a substantial fraction of their epitope is comprised of glycan (*32*). To investigate if a similar phenomenon occurred in the CAP256SU SHIV recalled lineages, we quantified the protein and glycan components of the respective epitopes for CH35-Apex1.08, Apex2.03, and Apex19.01 (Fig. 6E). While each CH35 lineage had approximately the same total interactive surface area (ranging from 1931 to 2164 Å^2^), CH35-Apex2.03 had a lower fraction mediated by glycan recognition – 0.23 compared to 0.52 and 0.50 for CH35-Apex1.08 and CH35-Apex19.01, respectively. An uncommon feature of the CH35-Apex2.03 mode of recognition that contributed to the glycan-deficient interface was the positioning of LCDR3 directly on top of Env residue 156, resulting in complete disordering of the N156 glycan (fig. S12D). These data suggest that limited glycan recognition can restrict the ability of high-affinity V2 apex-targeted antibodies to neutralize HIV strains, as is the case for CH35-Apex2.03.

The structural analyses thus far revealed two plausible explanations for the lack of neutralization by the CH35-Apex2 lineage, but it did not yet provide a molecular basis for the 2.5-fold difference in heterologous breadth observed between the CH35-Apex1 and CH35-Apex19 lineages. We therefore compared epitope residues in less-conserved regions within V1V2, such as the hypervariable V2 loop (Env positions 179-190) (*64*). CH35-Apex1.08 and CH35-Apex19.01 recognized hypervariable V2 from two protomers, although there was a much larger interface with this loop on protomer_B_ mediated by HCDR1 for both lineages (Fig. 6B). However, the HCDR1 of CH35-Apex19.01 was positioned closer to hypervariable V2 and induced conformational flexibility such that a stretch of three Env residues could not be accurately modeled, indicating an unstable or unfavorable interaction with this loop due in part to their close proximity to each other (Fig. 6F and fig. S12E). In addition, an Arg residue in the CH35-Apex19.01 HCDR3 had fully extended to form a salt bridge with Env residue E190; in contrast, CH35-Apex1.08 recognition was restricted to HCDR1. This structural insight suggests CH35-Apex19.01 to be more sensitive than CH35-Apex1.08 to the sequence and length diversity in hypervariable V2 among heterologous HIV strains, thereby restricting the neutralization breadth of the former.

Collectively, these structures show that bnAb function cannot be predicted from sequence signatures alone, as antibodies with canonical motifs and C-strand recognition may still adopt binding geometries incompatible with neutralization. High-resolution structural analyses were critical for distinguishing true bnAbs from “born-wrong” bnAb-like antibodies by revealing differences in angle of approach, glycan engagement, and epitope footprint that directly impact neutralization breadth.

## Discussion

This study provides mechanistic insight into how germline-targeting vaccination can initiate broadly neutralizing antibody (bnAb) responses against the HIV V2-apex. Our findings demonstrate that the quality of priming—particularly the recruitment of diverse bnAb precursor lineages and their early clonal expansion—strongly influences subsequent serum neutralization breadth. Rather than arising solely from rare stochastic events, broad serum neutralization was associated with the early activation of diverse long-CDRH3 precursors, followed by the preferential expansion and maturation of selected lineages.

Although Q23-APEX-GT2 consistently engaged V2-apex–directed B cells in all animals, serum neutralization emerged primarily in those that primed multiple diverse V2-apex long CDRH3 lineages and supported their robust early clonal expansion. Quantitative lineage tracking further revealed that serum neutralization was largely driven by one or two dominant clones whose expansion magnitude, combined with affinity maturation, predicted serum ID_50_ titers. These observations support a model in which early clonal amplification establishes a competitive advantage within germinal centers, increasing the probability that structurally favorable lineages re-enter iterative cycles of mutation and selection. Strong Tfh support (*67, 68*), efficient antigen capture (*69*), and sustained germinal center activity (*70, 71*), likely reinforce this process, enabling selected precursors to accumulate the mutations required to accommodate the dense glycan environment of the V2-apex and evolve toward breadth.

Importantly, our data suggest that successful priming must surpass a critical quantitative threshold: activating a single precursor is unlikely to be sufficient given the stringent structural constraints of bnAb development. Instead, recruiting multiple candidate lineages creates a diversified evolutionary landscape in which at least a subset can follow productive expansion and maturation trajectories. Robust activation and clonal expansion of diverse candidate lineages may promote effective trafficking of evolving B cells across lymph nodes throughout the lymphatic network, maximizing germinal center evolution and establishing sufficient memory. Their increased numbers also position these clones for efficient recall upon a polishing boost, amplifying responses, driving further maturation, and supporting long-lived plasma cell deposition for driving durable serum neutralizing antibody titers (fig. S13). As modeled here by CAP256.SU SHIV infection, exposure to a native-like Env appears to function as a stringent selection filter that preferentially amplifies those lineages with compatible angles of approach and epitope engagement, thereby accelerating the emergence of serum breadth.

A key conceptual advance of this work is the distinction between bnAb-associated sequence features and functional neutralization capacity. Several expanded lineages in CH35 monkey displayed canonical genetic signatures of V2-apex bnAbs—including long CDRH3 loops, anionic motifs, and characteristic D-gene usage—yet failed to acquire neutralizing activity despite extensive recall and somatic hypermutation. The CH35-Apex2 lineage exemplifies this disconnect, although it expanded robustly and exhibited cross-reactive binding to diverse tier 2 HIV Env trimers, it remained entirely non-neutralizing, suggesting an intrinsic structural incompatibility with productive epitope engagement needed for neutralization. These findings caution against relying solely on sequence-based metrics to define bnAb precursors and highlight the necessity of functional and structural validation to distinguish productive lineages from “dead-end” bnAb-like lineage responses.

Collectively, our results support a revised framework for germline-targeting vaccine design in which priming efficiency is evaluated not only by epitope specificity but also by the breadth of diverse precursor recruitment and the magnitude of their early expansion. Effective vaccines must seed a sufficiently large and diverse pool of candidate lineages while simultaneously guiding them toward favorable maturation solutions. The emergence of non-neutralizing yet highly competitive clones underscore the importance of immunogen designs that impose the correct selection pressures early in the response. Accordingly, priming immunogen design strategies should prioritize enhanced binding to a diverse range of bnAb precursors rather than a narrow subset (*32*), increased affinity and avidity for precursor lineages (*22, 72, 73*). Additional approaches, including T helper epitope conditioning (*74–76*), targeted delivery to lymphoid follicles to enhance antigen presentation (*77*), and controlled antigen-release technologies that sustain germinal center responses (*58, 59, 78–80*) may further improve precursor recruitment, expansion, and maturation. The benefits of these approaches, individually and in combination, warrant further investigation.

Our findings also raise the possibility that germline-targeting immunization and natural HIV-1 infection differ in the breadth of precursor recruitment and the stringency of subsequent selection. Native Env on infectious virions predominantly adopts a closed, prefusion trimer configuration, imposing structural constraints on B-cell recognition. Engineered germline-targeting immunogens, by contrast, are designed to engage otherwise rare bnAb precursors and may recruit a broader range of lineages, including those that cannot productively recognize native Env. This creates a potential trade-off: broader precursor recruitment may increase the probability of engaging lineages capable of developing breadth, but it may also generate expanded responses that ultimately prove nonproductive. Conversely, more stringent antigenic selection may favor structurally compatible lineages while limiting initial precursor recruitment. Determining how these competing pressures shape B-cell evolution during vaccination versus natural infection could inform strategies to improve both the efficiency and selectivity of bnAb induction.

More broadly, our findings help explain why bnAb induction has been historically rare: the pathway requires both quantitative success in precursor activation, and qualitative success in clonal trajectory. By demonstrating that targeted priming can establish *bona fide* bnAb precursors capable of rapid maturation upon appropriate boosting, this work provides a proof-of-principle that rational vaccination strategies can overcome key developmental barriers. Future efforts should therefore focus on optimizing immunogens and boosting regimens that preferentially expand structurally competent precursors while limiting the persistence of off-pathway clones, thereby increasing the reproducibility and predictability of bnAb induction.

## ACKNOWLEDGEMENTS

We thank the staff at Bioqual for exceptional care and assistance with nonhuman primates.

## Funding

This work was supported by National Institute of Allergy and Infectious Diseases National Institutes of Health grants R01 AI 167716 (R.A.), R61/R33 AI 161818 (R.A., G.M.S.), U19 AI 188562 (R.A.), P01 AI 177683 (R.A. and B.B.) R01 AI 165080 (G.M.S.) and R37 AI 150590 (B.H.H.). The data for this (manuscript or presentation) were generated in the Penn Cytomics and Cell Sorting Shared Resource Laboratory at the University of Pennsylvania (RRID:SCR_022376). Penn Cytomics is partially supported by the Abramson Cancer Center NCI Grant (P30 016520). The content is solely the responsibility of the authors and does not necessarily represent the official views of the National Institutes of Health. This study was made possible by the generous support of the Bill and Melinda Gates Foundation through the Collaboration for AIDS Vaccine Discovery (CAVD) grants INV041767 and INV064777 (G.M.S., and R.A.) We thank Carlos Diazgranados and Pervin Anklesaria from Gates Foundation for their support and helpful discussions.

## Author contributions

B.L., Y.Z., R.S.R., P.D.K, G.M.S and R.A. conceived and designed experiments. B.L., N.M, S.C., X.L., G.A. and C.M. prepared immunogens, isolated mAbs, performed binding assays. B.L., X.L. and C.L.M performed neutralization assays. B.L., Y.Z., Q.H., B.B. and A.B. performed B cell repertoire sequencing and analysis. B.L., Y.Z., Q.H., A.L.V. and G.G. performed single B cell sorting and B cell sequence analysis. B.L., X.L., G.G., L.T., R.R.C., P.O., K.A., Y.Z., S.A., T.V.S, A.S. and M.K. cloned, expressed, purified and tested monoclonal antibodies and trimers and probes for B cell sorting. M.G.L, A.N.S., F.B.R., B.H.H. and G.M.S. led NHP studies. D.J.I. synthesized and provided SMNP adjuvant. R.S.R., L.S., Z.S. and P.D.K. performed cryo-EM high-resolution structures of antibodies in complex with trimers and structural analysis. B.L., Y.Z., Q.H. and A.B. analyzed repertoire and bulk cell NGS data. B.L., Y.Z., Q.H., S.D.S. and A.W prepared the figure. B.L., Y.Z., R.S.R., D.R.B., P.D.K., G.M.S. and R.A. wrote the manuscript with input from all listed authors.

## Competing interests

R.A., G.M.S., R.N., X.L., B.L., R.R.C., Y.Z., K.A., N.M., S.C., G.A., D.R.B., and W.H. are listed as inventors on pending patent applications jointly filed by the University of Pennsylvania and Scripps Research, related to the immunogens used in this study. All other authors declare no competing interests.

## Data and materials availability

All data supporting the findings of this study are available within the paper and its Supplementary Information. Any additional information is available from the corresponding author upon reasonable request. Antibody sequences, 10x Genomics single-cell sequencing data, and bulk NGS datasets have been deposited in GenBank under accession numbers PZ018568 -PZ018870, BioProject PRJNA1427718, with BioSample accessions SAMN55930814-SAMN55930820, SAMN55932837-SAMN55932946 and associated SRA files (SRR37363789-SRR37363795, SRR37355277-SRR37355386). Negative-stain electron microscopy (ns-EM) maps of CH35-Apex1, CH35-Apex2.03 and CH35-Apex19.01 Fabs in complex with the Q23-APEX-GT2 protein have been deposited in the RCSB Protein Data Bank (RCSB PDB) under accession 11LA and 11KZ and the Electron Microscopy Data Bank (EMDB): EMD-75793 and EMD-75792.

## Supplementary Materials

### MATERIALS AND METHODS

#### Cell Lines

Expi293F cells (Gibco, Cat# A14527) were maintained in Expi293 Expression Medium (Gibco, Cat# A1435101) at 37°C under 8% CO₂ with agitation at 125 rpm. HEK293F cells (Gibco, Cat# A14527) were grown in Freestyle medium (Gibco, Cat# 12338-018) using the same temperature, CO₂, and shaking conditions. HEK293T cells (ATCC, Cat# CRL-3216) were cultured at 37°C in 5% CO₂ in Dulbecco’s Modified Eagle Medium (DMEM; Corning, Cat# 10-017-CV) supplemented with 10% heat-inactivated fetal bovine serum (FBS; Thermo Fisher, Cat# MT35016CV), 4 mM L-glutamine (Corning, Cat# 25-005-CI), and 1% penicillin-streptomycin (P/S; Corning, Cat# 30-002-CI). TZM-bl 931 cells (NIH AIDS Reagents Program) were used for pseudovirus neutralization assays as described previously.

#### Animal models

Six Indian rhesus macaques were used in this study in accordance with Association for Assessment and Accreditation of Laboratory Animal Care (AAALAC) guidelines. Animals were housed at Bioqual, Inc. (Rockville, MD), sedated for blood collection, and cared for under AAALAC standards and best-practice guidelines. All procedures were approved by the Institutional Animal Care and Use Committees (IACUC) of the University of Pennsylvania (protocol 807492) and Bioqual (protocol 24-072).

#### Stabilized Env Expression and Purification

All stabilized Env constructs were designed as previously described (*81*). HEK293F cells were transfected with plasmids encoding the indicated Env trimers using PEI-MAX 40K (Kyfora, Cat# 24765-1). Culture supernatants were harvested four days after transfection. Trimers were purified by affinity chromatography using either agarose-bound Galanthus nivalis lectin (GNL; Vector Labs, Cat# AL-1243-5) or TOYOPEARL AF-Tresyl-650M beads (TOSOH, Cat# 0014472) conjugated to the PGT145 broadly neutralizing antibody (bnAb). Eluates were further polished by size-exclusion chromatography (SEC) on a Superdex 200 Increase 10/300 GL column (GE Healthcare, Cat# GE28-9909-44) equilibrated in PBS. His-tagged proteins were purified using HisPur Ni-NTA resin (Thermo Fisher Scientific), buffer-exchanged into PBS, and subjected to an additional SEC step using a Superdex 200 column (Cytiva). SEC fractions corresponding to the trimer peak were pooled and used for ELISA and BLI binding studies, immunizations, and as baits for antigen- or epitope-specific single B cell sorting by flow cytometry.

#### Enzyme-Linked Immunosorbent Assay (ELISA)

ELISAs were performed following established protocols using biotinylated proteins captured on streptavidin-coated plates. 96-well half-area clear plates (Corning, Thermo Fisher Scientific) were coated overnight at 4°C with 2 μg/mL streptavidin (Jackson ImmunoResearch, Cat# 016-000-113) and washed three times with 1× PBS/0.05% Tween (Sigma-Aldrich, Cat# 1003620819). Plates were blocked for 1 h at room temperature (RT) with 3% BSA in PBS (Sigma-Aldrich, Cat# A9418-500G), then emptied and tapped dry. Biotinylated antigens (2 μg/mL) diluted in 1% BSA/1× PBS/0.05% Tween were added and incubated for 1.5 h at RT, followed by three washes. Serially diluted monoclonal antibodies (mAbs) or serum samples were added for 1.5 h at RT and washed. Alkaline phosphatase-conjugated secondary antibodies (Jackson ImmunoResearch, Cat# 109-055-170) were applied at 1:1000 (50 μL/well) for 1 h at RT. Phosphatase substrate (Thermo Fisher Scientific, Cat# S0942-200TAB) was prepared by dissolving one tablet per 5 mL alkaline staining buffer (1 L: 2.03 g MgCl₂·6H₂O, Fisher Bioreagents, Cat# BP214-500; 8.4 g Na₂CO₃, Sigma-Aldrich, Cat# S7795-500G; adjust pH to 9.8; bring to 1 L with MilliQ; add 1.0 g NaN₃, Sigma-Aldrich, Cat# S2002-100G; filter through 0.22 μm; store at 4°C). Absorbance at 405 nm was read after 20 min of development on a Synergy HTX multi-mode reader.

#### Biolayer Interferometry

Isolated mAbs was performed by BLI using 10 μg/mL IgG on an Octet Red96 (ForteBio). IgGs were loaded onto Protein A (ProA) biosensors (Sartorius, Cat# 18-5012) to a response of at least 1.0 nm, then equilibrated in running buffer (1× PBS, 0.02% Tween20, pH 7.4). Sensors were exposed to 500 nM trimer for 120 s (association) followed by 240 s in running buffer (dissociation). To measure polyclonal serum IgG responses, sera were diluted 1:50. Biotinylated trimers were captured onto streptavidin (SA) biosensors (Sartorius, Cat# 18-5020) to at least 1.0 nm on an Octet Red96, equilibrated in running buffer, then dipped into diluted serum for 120 s (association) and returned to running buffer for 240 s (dissociation). BLI Kd measurements were performed using Fab versions of mAbs. Fab heavy-chain plasmids were generated by introducing a His-Avi tag followed by a stop codon upstream of the Fc disulfide bond. For Fab biotinylation, heavy- and light-chain plasmids were co-transfected with BirA into Expi293 cells (Thermo Fisher Scientific, Cat# A14527) at a 2:2:1 plasmid ratio (HC:LC:BirA) using FectoPRO (Polyplus, Cat# 116-001). At 24 h post-transfection, cultures were supplemented with 0.3 M valproic acid (Sigma, Cat# P4543-100G) and 40% glucose (Gibco, Cat# A2494001). Fab fragments were purified from culture supernatants four days post-transfection using Ni Sepharose 6 Fast Flow (Cytiva, Cat# 17531802). Eluted proteins were buffer-exchanged into PBS and concentrated with 10 kDa MWCO centrifugal filters (Millipore, Cat# UFC905024), then further purified by SEC on a Superdex 200 Increase 10/300 GL column (Sigma-Aldrich, Cat# GE28-9909-44). Peak fractions were pooled and concentrated. For kinetic runs, 10 μg/mL Fab was captured on SA biosensors (Sartorius, Cat# 18-5020) to at least 1.0 nm, equilibrated in running buffer, and exposed to a two-fold serial dilution of trimers starting at 500 nM for 120 s (association), followed by 240 s dissociation in running buffer.

#### Neutralization Assay

Sera (starting dilution 1:20) or monoclonal antibodies (pre-infection mAbs starting concentration 300 μg/mL, post-infection 30ug/mL) were serially diluted three-fold in 25 μL complete DMEM and incubated with 25 μL HIV-1 Env-pseudotyped virus for 60 min at 37°C in duplicate 96-well culture plates. TZM-bl cells (20,000 cells/well) in the presence of 20 μg/mL DEAE-Dextran were added (50 μL) and plates were incubated (*82*). Control wells contained cells only (background) or virus only (maximal entry). Luciferase activity was measured at 72 h using the Bright-Glo Luciferase Reporter Assay (Promega, Cat# E2650) on a Synergy HTX multi-mode luminometer. Percent neutralization was calculated as ((RLU_virus − RLU_test) / RLU_virus) × 100 after subtraction of background RLU from uninfected wells. Neutralizing serum titers (ID50) and antibody IC50 values were determined by fitting a four-parameter nonlinear dose-response inhibition curve.

#### Neutralization Contribution Score (NCS)

For each animal, the contribution of each V2-apex lineage was estimated by integrating lineage expansion measured by bulk BCR sequencing and the neutralization potency of the corresponding monoclonal antibody.

For lineage i,

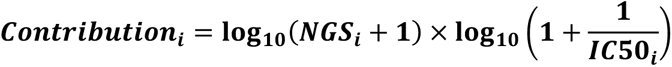

Where ***NGS_i_***is the bulk sequencing abundance of lineage i at the indicated time point. ***IC*50*_i_*** is the neutralization IC50 (µg/mL) of the corresponding monoclonal antibody against the matched virus.

The Neutralization Contribution Score (NCS) for each animal was calculated as

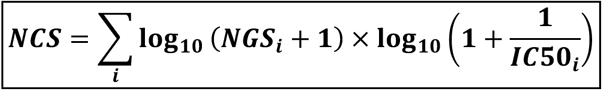

No parameter fitting or regression optimization was performed.

Associations between NCS and serum neutralization ID50 were evaluated using both Pearson and Spearman correlation analyses.

#### Immunization in rhesus macaques, blood and lymph node processing

Six rhesus macaques (3-5 years old; balanced by sex) were immunized with the germline-targeting trimer Q23-APEX-GT2 formulated with SMNP adjuvant (*57*), and the prime (week 0) was given as escalating dose (DE) over 2 weeks (*58, 59*), followed by bolus boost at week 10, using as previously described^13^. Animals were challenged at week 18 with CAP256.SU SHIV. Peripheral blood was collected into sterile vacutainers (BD Vacutainer, Cat# 364606) containing acid citrate dextrose formula A (ACD-A). To process, 40 mL ACD-A blood was transferred to a sterile 50 mL conical tube and centrifuged at 1000g for 10 min at 20°C. Plasma was removed without disturbing the buffy coat or red cell pellet and clarified at 1500g for 15 min at 20°C. Cell-depleted plasma was aliquoted into 1 mL cryovials (Sarstedt, Cat# 72.694.396) and stored at - 80°C. The remaining cell fraction was resuspended in an equal volume of HBSS without Ca²⁺/Mg²⁺ (HBSS−/−; Gibco, Cat# 14175-079) containing 2 mM EDTA (Invitrogen, Cat# 15575-00). Suspensions were split across four 50 mL conical tubes, brought to 35 mL with HBSS−/− + EDTA, and underlaid with 14 mL 96% Ficoll-Paque Plus (Cytiva, Cat# 17144003). Gradients were centrifuged at 725g for 20 min at 20°C with slow acceleration/deceleration. Mononuclear cells at the interface were collected, washed in HBSS−/− + EDTA (200g, 15 min, 20°C), and resuspended in 40 mL HBSS with Ca²⁺/Mg²⁺ (HBSS+/+; Gibco, Cat# 24020-117) supplemented with 1% FBS (Cytiva, Cat# SH300.71.03). Cells were centrifuged at 200g for 15 min at 20°C to pellet white blood cells while leaving platelets in suspension; supernatant was removed and pellets were gently resuspended and brought to 10 mL with HBSS+/+ + 1% FBS. Cell counts and viability were determined using ViaStain AOPI solution (Revvity, Cat# CS2-0106-25ml) on a Cellometer Auto 2000 (Revvity, Waltham, MA). Cells were then pelleted (300g, 10 min, 20°C), resuspended in CryoStor CS5 (Stemcell Technologies, Cat# 07930) at 5-10 × 10⁶ cells/mL, aliquoted into 1 mL cryovials (Thermo Scientific, Cat# 374503), and frozen overnight at -80°C in a CoolCell LX (Corning, Cat# 432002) or FTS30 (Corning, Cat# 432006) container before transfer to vapor-phase liquid nitrogen. Mononuclear cells from lymph nodes (LNs) were processed similarly. Immediately after axillary and/or inguinal LN excision, tissues were placed in RPMI1640 (Corning, Cat# MT15040CV) on wet ice and processed within 6 h. Nodes were minced with a sterile scalpel, passed through a sterile cell strainer (Falcon, Cat# 352360), and the resulting cell suspension was subjected to Ficoll density-gradient purification as above, substituting RPMI1640 containing 10% FBS for HBSS.

#### Flow cytometry staining

For B and T cell profiling, multiparameter flow cytometry was performed using specific cell surface markers, as previously described (*62*). Cryopreserved PBMC and lymph node (LN) samples were thawed in RPMI 1640 (Thermo Fisher Scientific, Cat# MT15040CV) supplemented with 50% fetal bovine serum (FBS). Cells were washed in FACS buffer (PBS containing 2% FBS and 2 mM EDTA; Invitrogen, Cat# 15575-038). Fluorescent antigen probes were generated by stepwise addition of fluorophore-conjugated streptavidin to biotinylated Q23-APEX-GT2, C1080-SCT1, CAP256.SU-SCT9, and the epitope knockout probe Q23-APEX-GT2 dKO over 45 min at room temperature in PBS. Cells were incubated with antigen probes for 30 min at 4 °C. Cells were first washed once with PBS and stained with a LIVE/DEAD viability dye (eFluor506; Thermo Fisher Scientific, 65-0866-18, 1:500) for 20 min at room temperature. Following washing, cells were stained with a master mix of surface antibodies including: CD8α (RPA-T8, APC-eFluor780, Thermo Fisher Scientific, 47-0088-42, 1:100), CD16 (CB16, APC-eFluor780, Thermo Fisher Scientific, 47-0168-42, 1:100), CD14 (M5E2, APC-Cy7, BioLegend, 301820, 1:100), CD20 (2H7, PerCP-Cy5.5, BioLegend, 302326, 1:100), IgG (G18-145, BUV737, BD Biosciences, 612819, 1:100), IgM (MHM-88, Pacific Blue, BioLegend, 314514, 1:50), IgD (polyclonal, PE, Southern Biotech, 2030-09, 1:50), CD38 (OKT10, PE-Cy5, Caprico Biotechnologies, 100875, 1:50), CD71 (L01.1, BUV615, BD Biosciences, 751091, 1:20), CD27 (O323, BV785, BioLegend, 302832, 1:50), CD3 (SP-34, BUV395, BD Biosciences, 564117, 1:40), CD4 (OKT4, BUV496, BD Biosciences, 750980, 1:100), PD-1 (EH12.2H7, BV711, BioLegend, 329928, 1:20), and CXCR5 (Mu5UBEE, PE-Cy7, Thermo Fisher Scientific, 25-9185-42, 1:20). Samples were acquired on a BD FACSDiscover A8 Research Cell Analyzer (Penn Cytomics and Cell Sorting Shared Resource Laboratory, University of Pennsylvania) and analyzed using FlowJo (BD Biosciences).

#### Flow Cytometry Antigen-Specific B Cell Sorting

Cryopreserved LN and PBMC samples were thawed and resuspended in pre-warmed medium consisting of 50% RPMI (Thermo Fisher, Cat# MT15040CV) and 50% FBS. PBMC samples were treated individually with ACK lysing buffer (Gibco, Cat# A10492-01) to remove residual red blood cells. LN and PBMC cells were washed in FACS buffer (PBS with 2% FBS and 2 mM EDTA; Invitrogen, Cat# 15575-038) and viability was assessed using trypan blue (Invitrogen, Cat# T10282) on a Countess cell counter. Cells were stained with fluorophore-conjugated antigens on ice for 30 min in the dark. Avi-tagged, biotinylated Q23-APEX-GT2 trimer and variants (dKO, 169E-171E) were conjugated to streptavidin fluorophores BV421 (BD Horizon, Cat# 563259), AF647 (Invitrogen, Cat# S21374), and AF488 (Invitrogen, Cat# S11223) at RT for 30 min in the dark prior to staining. Surface antibodies to CD3 (BD Pharmingen, Cat# 557757), CD4 (BioLegend, Cat# 317418), CD8 (BioLegend, Cat# 557760), CD14 (BD Pharmingen, Cat# 561384), CD19 (BioLegend, Cat# 302230), CD20 (BioLegend, Cat# 302326), IgG (BD Horizon, Cat# 564230), and IgM (BioLegend, Cat# 314508) were added and incubated for 30 min on ice in the dark. FVS510 Live/Dead stain (Thermo Fisher Scientific, Cat# L34966) was applied at 1:250 for 15 min on ice in the dark. Cells were washed, passed through a cell strainer into 5 mL round-bottom tubes (Corning, Cat# 352058), and sorted on a BD FACSMelody into 96-well PCR plates as single cells. RT-PCR was performed using SuperScript IV with IgH, IgK, and IgL gene-specific primers. Paired heavy and light chains were amplified by PCR and visualized on 2% E-gel 96 agarose gels with SYBR Safe (Thermo Fisher Scientific, Cat# G720802). Wells with successful amplification products were sequenced by Sanger sequencing (Azenta).

#### Antigen-specific B Cell lineage analysis using 10x Genomic feature barcoding method

Antigen- and epitope-specific B cells were isolated by flow cytometry, followed by recovery of paired antibody lineage sequences using the 10x Genomics feature barcoding platform, as previously described (*83*). A well of a 96-well PCR plate was pre-coated with FBS to cushion cells during sorting and the FBS was removed immediately before sorting. Cryopreserved LN and PBMC samples were thawed and resuspended in pre-warmed 50% RPMI/50% FBS. PBMCs were treated individually with ACK lysing buffer to remove residual red blood cells. Cells were washed with 10% FBS and viability was assessed by trypan blue on a Countess counter. LN and PBMC cells were stained with conjugated antigens for 30 min on ice in the dark. Conjugated antigens were prepared by incubating Avi-tagged, biotinylated Q23-APEX-GT2 trimer and variants (dKO, 169E-171E) with streptavidin fluorophores at RT for 30 min in the dark. Cells were then stained for 30 min on ice in the dark with antibodies to CD3, CD4, CD8, CD14, CD19, CD20, IgG, and IgM, along with anti-human hashtag antibodies (BioLegend, TotalSeq-C). FVS510 Live/Dead stain was added at 1:250 for 15 min on ice in the dark. Cells were washed, filtered into 5 mL round-bottom tubes, and sorted on a BD FACSMelody into the pre-coated well. Single-cell processing was performed using Chromium GEM-X Single Cell 5′ Reagent Kits v3 with feature barcode technology for cell surface protein (10x Genomics, CG000734, Rev A) to generate GEMs, amplify cDNA, perform V(D)J amplification, and construct libraries for V(D)J, gene expression, and cell surface protein profiling. Libraries were sequenced on an Illumina NextSeq 1000/2000 platform and initial processing used 10x Cloud Analysis. Sequencing reads were further processed with barcounter v1.0.0 (https://doi.org/10.1101/2020.09.04.283887) and demultiplexed via deMULTIplex2 into V(D)J repertoire libraries (*84*).

#### Antibody Cloning

IgG heavy- and light-chain sequences derived from sorted B cells and exhibiting features characteristic of rhesus V2-apex broadly neutralizing antibodies were inserted into IgG1, Igκ, or Igλ AbVec expression vectors upstream of the corresponding constant regions. Inserts were introduced using AgeI/NheI (heavy chain), AgeI/BsiWI (κ light chain), or AgeI/SaII (λ light chain) restriction sites. Assembly was performed with NEBuilder HiFi DNA Assembly Master Mix (New England Biolabs, Cat# E5520S) according to the manufacturer’s instructions.

#### Monoclonal Antibody Expression and Purification

Paired heavy- and light-chain plasmids encoding rhesus or human V2-apex UCAs or monoclonal antibodies from immunized animals were co-transfected into Expi293 cells (Thermo Fisher Scientific, Cat# A14527) at a 1:1 ratio using FectoPRO (Polyplus, Cat# 116-001) per the manufacturer’s instructions. At 24 h post-transfection, cultures were supplemented with 0.3 M valproic acid (Sigma, Cat# P4543-100G) and 40% glucose (Gibco, Cat# A2494001) to enhance expression. Monoclonal IgGs were harvested from supernatants four days after transfection and purified using a 1:1 mixture of Protein A (Cytiva, Cat# 17127903) and Protein G Sepharose (Cytiva, Cat# 17061805). Bound antibodies were eluted with IgG elution buffer (Thermo Fisher Scientific, Cat# PI21009) and buffer-exchanged into PBS using 50 kDa MWCO centrifugal filters (Millipore, Cat# UFC905024).

#### Bulk Repertoire Sequencing

Cryopreserved LN and PBMC samples were thawed at 37°C and immediately transferred into pre-warmed 50% RPMI (Thermo Fisher, Cat# MT15040CV) / 50% FBS. Cells were pelleted and supernatants removed. For RNA extraction, pellets were lysed in RLT Plus buffer (Qiagen, Cat# 1053393) containing β-mercaptoethanol. Total RNA was purified with the RNeasy Plus Mini Kit (Qiagen, Cat# 74134), including on-column DNase I treatment (Qiagen, Cat# 79254). RNA quantity and quality were assessed by NanoDrop (Thermo Fisher Scientific) and Bioanalyzer (Agilent 2100) RIN score; only samples with RIN ≥ 7 were advanced. RT-PCR was performed using IgG- and IgM-specific primers. First-strand cDNA synthesis used SuperScript IV (Thermo Fisher Scientific, Cat# 1750150). cDNA was amplified by PCR and treated with ExoSAP-IT (Thermo Fisher Scientific, Cat# 78205). Heavy-chain V(D)J regions were amplified using a two-step PCR strategy with HotStarTaq Plus DNA Polymerase (Qiagen, Cat# 203603), beginning with framework primers and followed by nested amplification incorporating overhangs. Illumina adapters with unique dual indexes were added in the second PCR. Final libraries were size-selected and purified with SPRIselect beads (Beckman Coulter Genomics, Cat# B23318) at 0.8× bead-to-sample ratio to remove adapter dimers and short fragments. Library concentration was measured by Qubit (Thermo Fisher Scientific) and fragment size was verified using an Agilent 2100 Bioanalyzer High Sensitivity DNA chip. Libraries were pooled and sequenced on an Illumina NextSeq 1000/2000 platform.

#### B-cell Annotation / V(D)J Repertoire Analysis

Merged bulk sequencing reads were analyzed using KIMDB 1.1 (http://kimdb.gkhlab.se/) as the heavy-chain germline reference and the Ramesh et al. allele database for light-chain genes. B cell receptor sequences were filtered, quality controlled, merged, annotated, and deduplicated using abstar v0.7.3 (https://github.com/brineylab/abstar), AbUtils v0.5.0 (https://github.com/brineylab/abutils), and IgDiscover v1.0.5 to generate individualized immunoglobulin repertoire libraries for each rhesus macaque.

#### Antibody Clonotypes

Antibody clonotypes were defined by shared VH and JH gene usage, identical CDRH3 amino acid length, and near-identical CDRH3 AA sequences. Sequences with no CDRH3 mismatches were grouped into a single clonotype to reduce errors arising from amplification or sequencing.

#### Lineage Analysis and Tracing

Lineages were defined as collections of clonotypes sharing the same VH and JH gene assignments, identical CDRH3 lengths, and similar CDRH3 amino acid sequences. A CDRH3 amino acid similarity cutoff of 0.8 was applied to assign lineage membership, followed by manual inspection to reduce misclassification or omission. Where paired heavy- and light-chain sequences were available, lineage assignment was further supported by consistent light-chain V(D)J features. This criterion was applied to sequences generated by 10x Genomics sequencing and single B-cell sorting, for which both immunoglobulin heavy- and light-chain genes were sequenced to enable paired antibody characterization and monoclonal antibody synthesis. For these datasets, sequences were assigned to the same lineage based on consistency of heavy-chain features, including identical VH and JH gene usage, identical CDR3 length, and ≥80% amino acid sequence identity in the CDR3 region, together with consistent light-chain features, including identical VL/VK and JL/JK gene usage, identical CDR3 length, and similar CDR3 amino acid sequences. For the NGS dataset, light-chain sequences were not generated. Therefore, lineage assignment in NGS dataset was based solely on heavy-chain V(D)J features using the criteria described above. Lineage tracing applied these criteria across time points and cell types for each animal, integrating 10x Genomics and bulk NGS datasets.

#### Plasma vRNA quantification

Plasma viral load measurements were performed by the NIH/NIAID-sponsored Nonhuman Primate Virology Core Laboratory at the Duke Human Vaccine Institute, as previously described (*42*). Briefly, viral RNA was extracted from plasma and quantified by real-time quantitative PCR (qPCR) using automated QIAsymphony SP and QIAgility platforms (QIAGEN) on a QuantStudio 3 instrument (Thermo Fisher Scientific). Viral RNA copies per milliliter (cp/mL) were calculated from quantification cycle (Cq) values after quality control of positive and negative controls. The lower limit of quantification (LLOQ) ranged from 62 to 250 RNA copies/mL over the course of this study.

#### Viral sequencing

Single-genome sequencing of SHIV 3′ half-genomes was performed as previously described (*43, 85*). Briefly, viral RNA was extracted from plasma using the QIAamp Viral RNA Kit (Qiagen) and reverse transcribed using SuperScript III Reverse Transcriptase (Invitrogen). Viral cDNA was then endpoint diluted and amplified by nested PCR using primers and conditions described previously (PMID: 40502159). Geneious Prime was used for sequence alignment and downstream sequence analysis.

#### Monoclonal Fab preparation for Cryo-EM structural studies

Fab constructs were generated by introducing a C-terminal His-Avi tag and stop codon into the IgG heavy-chain plasmid upstream of the Fc disulfide-forming region. Truncated heavy- and light-chain vectors were co-transfected into Expi293 cells (Thermo Fisher Scientific, Cat# A14527) at equimolar ratios using FectoPRO (Polyplus, Cat# 116-001). At 24 h, cultures received valproic acid (0.3 M final; Sigma, Cat# P4543-100G) and glucose (40%; Gibco, Cat# A2494001). Supernatants were collected four days post-transfection and Fabs were purified using Ni Sepharose 6 Fast Flow resin (Cytiva, Cat# 17531802). Eluted proteins were exchanged into PBS and concentrated using 10 kDa MWCO centrifugal filters (Millipore, Cat# UFC905024), then further purified by SEC on a Superdex 200 Increase 10/300 GL column (Sigma-Aldrich, Cat# GE28-9909-44). Monomeric Fab peak fractions were pooled and concentrated for subsequent use.

#### Cryo-EM data collection and processing

The high-resolution structures Q23-APEX-GT2 envelope trimer bound by Fabs CH35-Apex2.03 and Apex19.01 were determined using single-particle cryo-EM as previously described (*60*). Samples were prepared by diluting purified Q23-APEX-GT2 to 2 mg/mL with a 3-fold molar excess of Fab:trimer and 0.005% (w/v) n-Dodecyl β-D-maltoside (DDM). Diluted complex samples were incubated on ice for 30 minutes. Freshly glow-discharged Copper C-flat Holey carbon-coated grids (CF-1.2/1.3 300 mesh; EMS) were applied with 3 uL of sample for 30 seconds in a Vitrobot Mark IV at room temperature with 100% humidity. Sample grids were blotted for 3 seconds and vitrified using liquid ethane. Cryo-EM data were collected on a FEI Titan Krios 300 kV cryo-transmission electron microscope equipped with a Gatan K3 detector operating in counting mode using Leginon (*86*) for the CH35-Apex19.01 dataset and using EPU (*87*) for the CH35-Apex2.03 dataset. The total electron dose of 58 e^-^/Å^2^ was fractionated over 50 raw frames. Defocus values were set to cycle between -0.80 and -2.0 μm. All data processing was done in cryoSPARC v3.2 (*88*). Particle picking was performed with unbiased blob picker. All homogenous and non-uniform 3D refinements of Fab-bound complexes were performed using C1 symmetry. Data acquisition and processing details for all structures are provided in Supplementary Table S6.

#### Atomic model building and refinement

Each atomic model was solved by iterative manual rebuilding in Coot (*89*) and real-space refinement in Phenix (*90*). Structure quality was periodically assessed using MolProbity (*91*) and EMRinger (*92*). The initial model used for the Q23-APEX-GT2 envelope trimer in the two structures presented here was our previously published apo Q23-APEX-GT2 structure (PDB-9nvv), which was fit into each respective cryo-EM reconstruction density using UCSF ChimeraX (*93*). The initial model of each Fab variable region (Fv) was first obtained using the AlphaFold3 server (*94*) and fit into the respective cryo-EM 3D reconstruction using ChimeraX followed by iterative refinement. Protein and N-linked glycan interface calculations were performed using PDBePISA (*95*). Final model statistics and validations are provided in Supplementary Table S6 and Extended Data Fig. 10.

#### Statistical Analysis

Statistical analyses were performed in GraphPad Prism and R. Two-tailed Student’s t-tests were used to assess significance unless otherwise stated. P values < 0.05 were considered significant and are denoted as: *P < 0.05; **P < 0.01; ***P < 0.001; ****P < 0.0001; ns, not significant. One-sided tests for positive Pearson correlation were computed to assess associations, with corresponding P values reported. Correlation analyses were performed using Pearson correlation coefficients. For analyses examining associations between B cell responses and serum antibody binding or neutralization, and between Tfh and GC B cell responses, one-sided tests for positive correlation were used because the analyses were based on a priori biological hypotheses predicting positive associations. For analyses examining the relationship between serum neutralization and monoclonal antibody IC50 values or NGS-derived measurements, two-sided tests were used because the direction and magnitude of the association were not prespecified. For simple linear regression analyses, statistical significance of the association was also assessed using a two-sided test of the regression slope.

## SUPPLEMENTARY FIGURES AND LEGENDS

**fig. S1.**
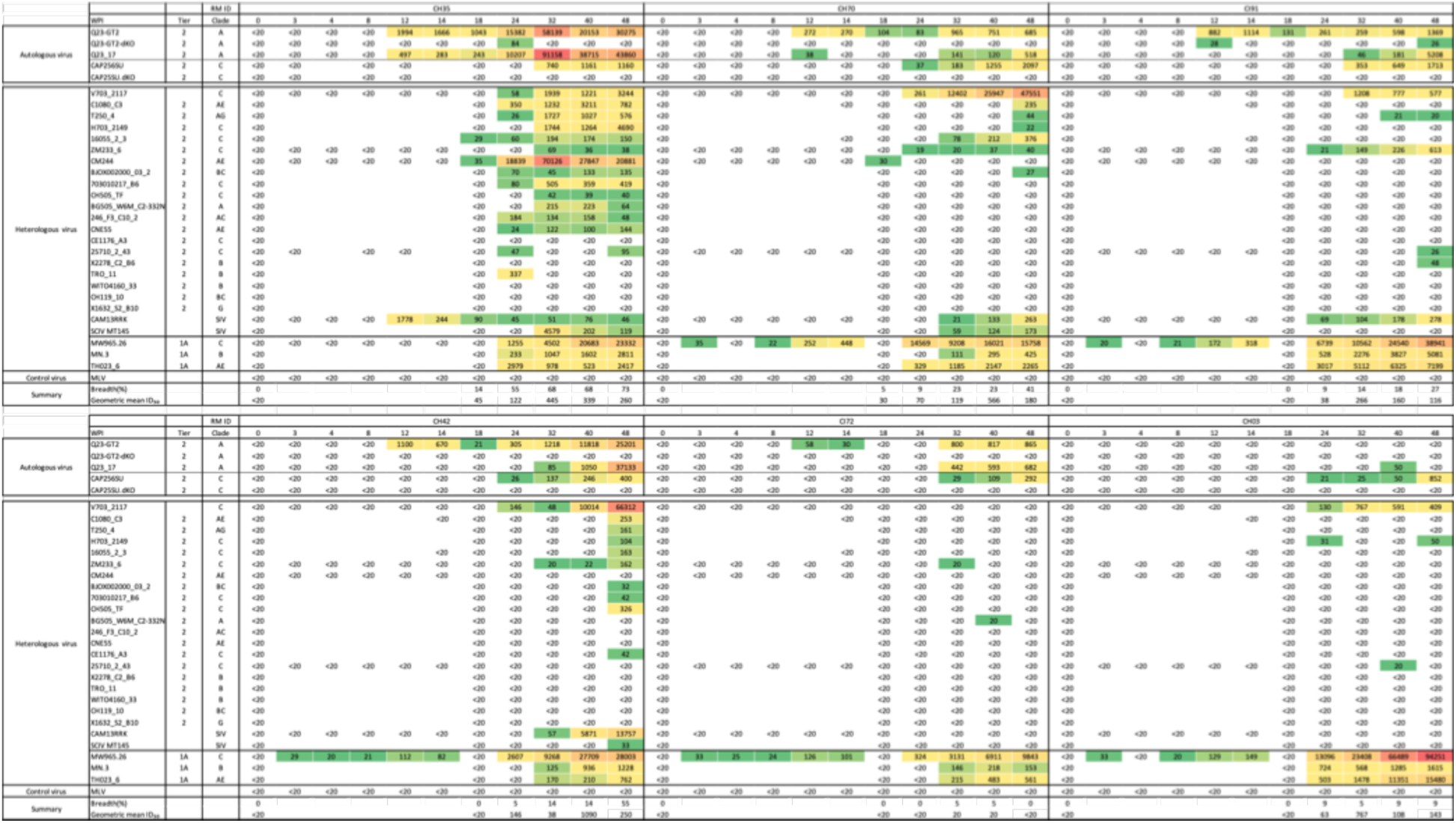
Longitudinal serum neutralization ID50 titers before and after CAP256.SU SHIV infection. Table summarizing serum neutralization ID50 titers measured at multiple time points for all animals before and after CAP256.SU SHIV infection. Sera were tested against a panel of HIV-1 viruses representing diverse clades, and ID_50_ values for each virus–time point combination are reported. Comparison of pre- and post-infection samples reveals an overall increase in neutralization breadth and potency following CAP256.SU infection, most prominently in animal CH35.

**fig. S2.**
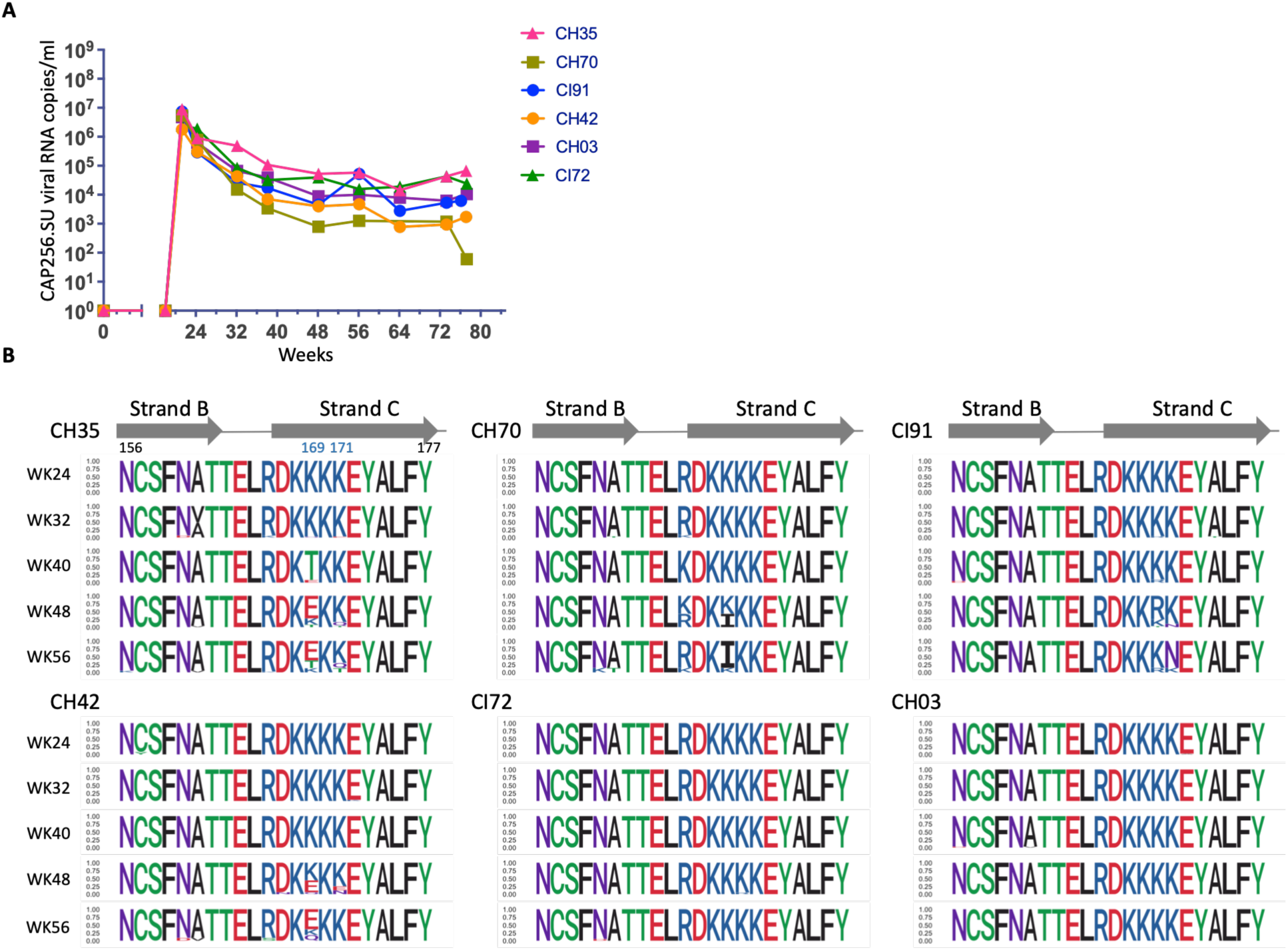
Evolution of CAP256.SU following SHIV infection. (**A**) Longitudinal plasma viral RNA load (copies/ml) in SHIV-infected rhesus macaques (**B**) Amino acid sequence alignment of strands B and C of the CAP256.SU Env V2 region across longitudinal timepoints following SHIV infection in all animals. Key residues within the core V2 epitope are highlighted. Positively charged amino acids are shown in blue, and negatively charged residues in red. The height of each residue at a given position reflects its mutation frequency at the respective timepoint. Escape-associated substitutions were observed at positions 169 and/or 171 in multiple animals, including K169E/K171Q-T in CH35, K169I in CH70, K171N in CI91, and K169E/K171E in CH42, resulting in reduced positive charge within the V2-apex core epitope.

**fig. S3.**
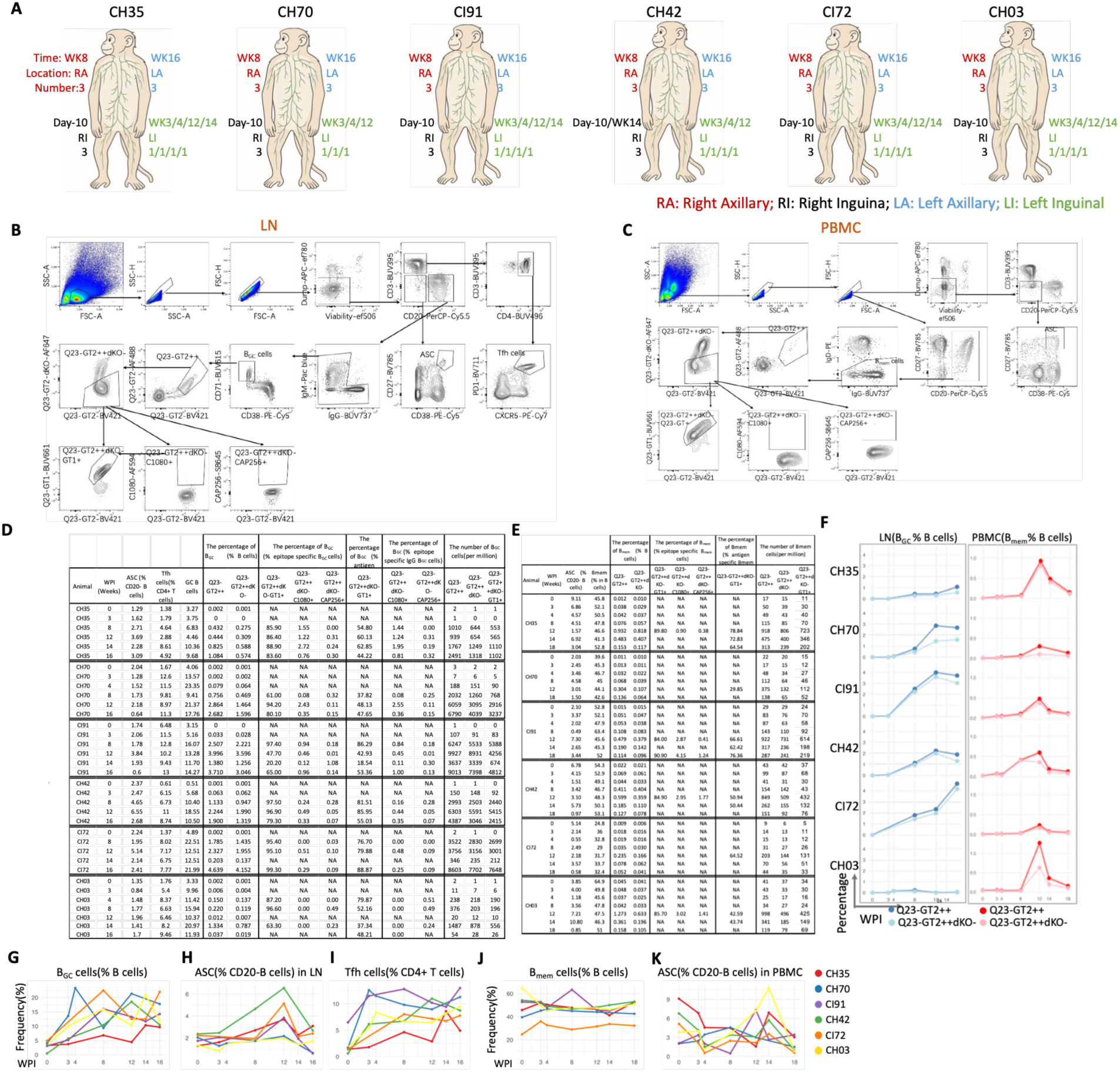
Flow cytometry analysis of B and T cell responses. (**A**) Diagram showing the anatomical locations of lymph node biopsies performed in all animals. (**B**) Gating strategy for lymph node (LN) analysis, including T follicular helper (Tfh), antibody-secreting cell (ASC), and germinal center B cell (B_GC_) populations. Gating for antigen-specific, epitope-specific, and heterologous trimer (Q23-GT1, C1080, and CAP256.SU)-specific B cells is shown. (**C**) Gating strategy for ASCs, memory B cells (B_mem_), and antigen-, epitope-, and heterologous trimer (Q23-GT1, C1080, and CAP256.SU)-specific B cells in peripheral blood mononuclear cells (PBMCs). (**D**) Summary table of all LN cell subsets analyzed by flow cytometry. (**E**) Summary table of all PBMC cell subsets analyzed by flow cytometry. (**F**) Line plots showing longitudinal frequencies of antigen-specific and epitope-specific B cells: GC B cells in LNs (left) and memory B cells in PBMCs (right), expressed as a percentage of total B cells. (**G**) Longitudinal frequency of B_GC_ (CD71⁺CD38⁻) as a percentage of total B cells (CD20⁺) in LNs. (**H**) Longitudinal frequency of ASCs (CD27⁺CD38⁺) as a percentage of total B cells in LNs. (**I**) Longitudinal frequency of Tfh cells (PD-1⁺CXCR5⁺) as a percentage of total CD4⁺ T cells in LNs. (J) Longitudinal frequency of B_mem_ (CD20⁺IgD⁻) as a percentage of total B cells in PBMCs. (K) Longitudinal frequency of ASCs (CD27⁺CD38⁺) as a percentage of total B cells in PBMCs.

**fig. S4.**
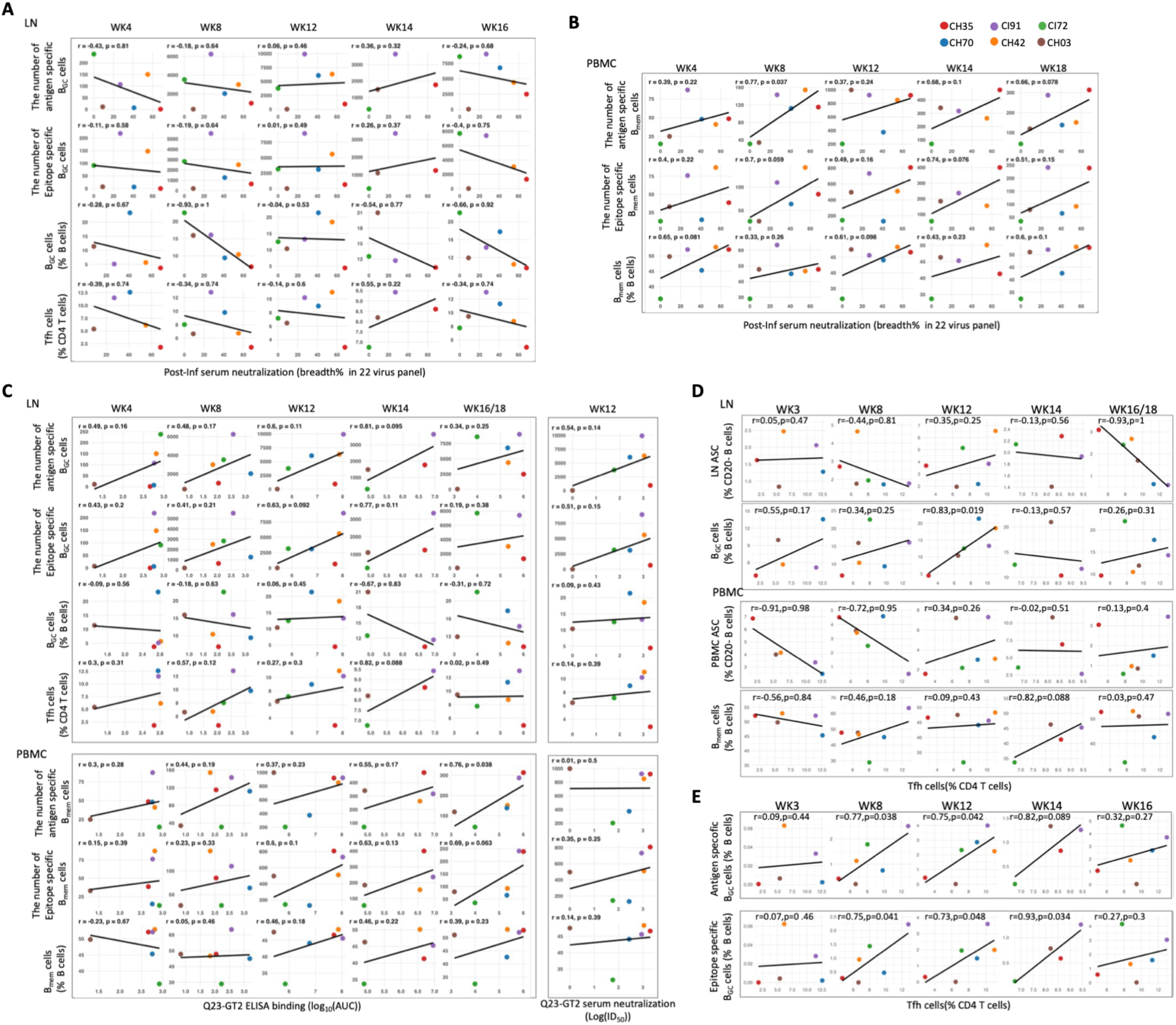
Correlation analyses between Tfh cells, B cell subsets, and serum antibody responses. **(A** and **B)** Correlation analyses between post-infection serum neutralization breadth and B cell populations in lymph nodes (LNs; A) and peripheral blood mononuclear cells (PBMCs; B) across longitudinal sampling time points. LN parameters include the numbers of antigen-specific and epitope-specific germinal center B cells (B_GC_) per million cells, the percentage of total B_GC_ cells, and the percentage of T follicular helper (Tfh) cells. PBMC parameters include the numbers of antigen-specific and epitope-specific memory B cells (B_mem_) per million cells and the percentage of total B_mem_ cells. (**C**) Correlation analyses of serum ELISA binding (log₁₀(AUC); left) and week 12 serum neutralization titers (ID₅₀) against Q23-APEX-GT2 (right) with B cell populations in lymph nodes (LNs; upper) and peripheral blood mononuclear cells (PBMCs; lower) across longitudinal sampling time points. LN parameters include the numbers of antigen-specific and epitope-specific germinal center B cells (B_GC_) per million cells, the percentage of total B_GC_ cells, and the percentage of T follicular helper (Tfh) cells. PBMC parameters include the numbers of antigen-specific and epitope-specific memory B cells (B_mem_) per million cells and the percentage of total B_mem_ cells. (**D**) Correlation between the percentage of T follicular helper (Tfh) cells and the frequencies of antibody-secreting cells (ASCs) and germinal center B cells (B_GC_) in lymph nodes (LNs; upper), or ASCs and memory B cells (B_mem_) in peripheral blood mononuclear cells (PBMCs; lower), across longitudinal sampling time points.. (**E**) Correlation between the frequency of antigen specific (Q23-APEX-GT2⁺⁺) B_GC_ cells and epitope specific (Q23-APEX-GT2⁺⁺dKO⁻) B_GC_ cells in LNs and the percentage of Tfh cells over time. Notably, Tfh cell frequencies showed strong correlations with antigen- and epitope-specific GC B cell responses

**fig. S5.**
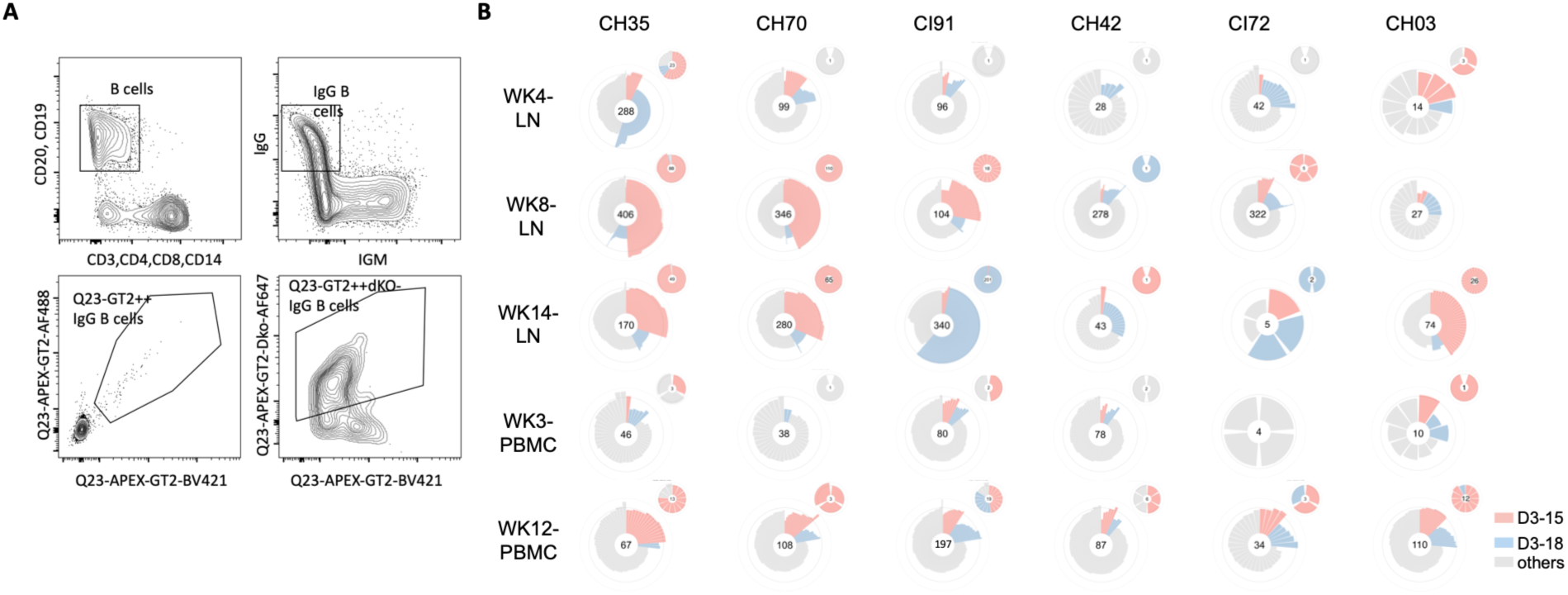
Characteristics of epitope-specific IgG⁺ B cell repertoires in Q23-APEX-GT2– immunized rhesus macaques.. (**A**) Representative flow cytometry plots illustrating the gating and single-cell sorting strategy used to isolate epitope-specific B cells (**B**) Sorted epitope-specific IgG⁺ B cells were isolated from weeks 4, 8 and 14 LNs and weeks 3 and 12 PBMCs of Q23-APEX-GT2–immunized rhesus macaques (RMs), followed by 10x Genomics sequencing. Circular bar plots illustrate D-gene usage among V2-apex–specific B cells. Key IGHD germline genes (IGHD3-15 and IGHD3-18) are individually highlighted, whereas all other D genes are grouped as “Others.” The dashed inner ring indicates the 22-amino acid threshold for CDRH3 length. Values in the center denote the total number of analyzed sequences, and smaller circular bar plots highlight sequences with ≥22 amino acid long CDRH3s.

**fig. S6.**
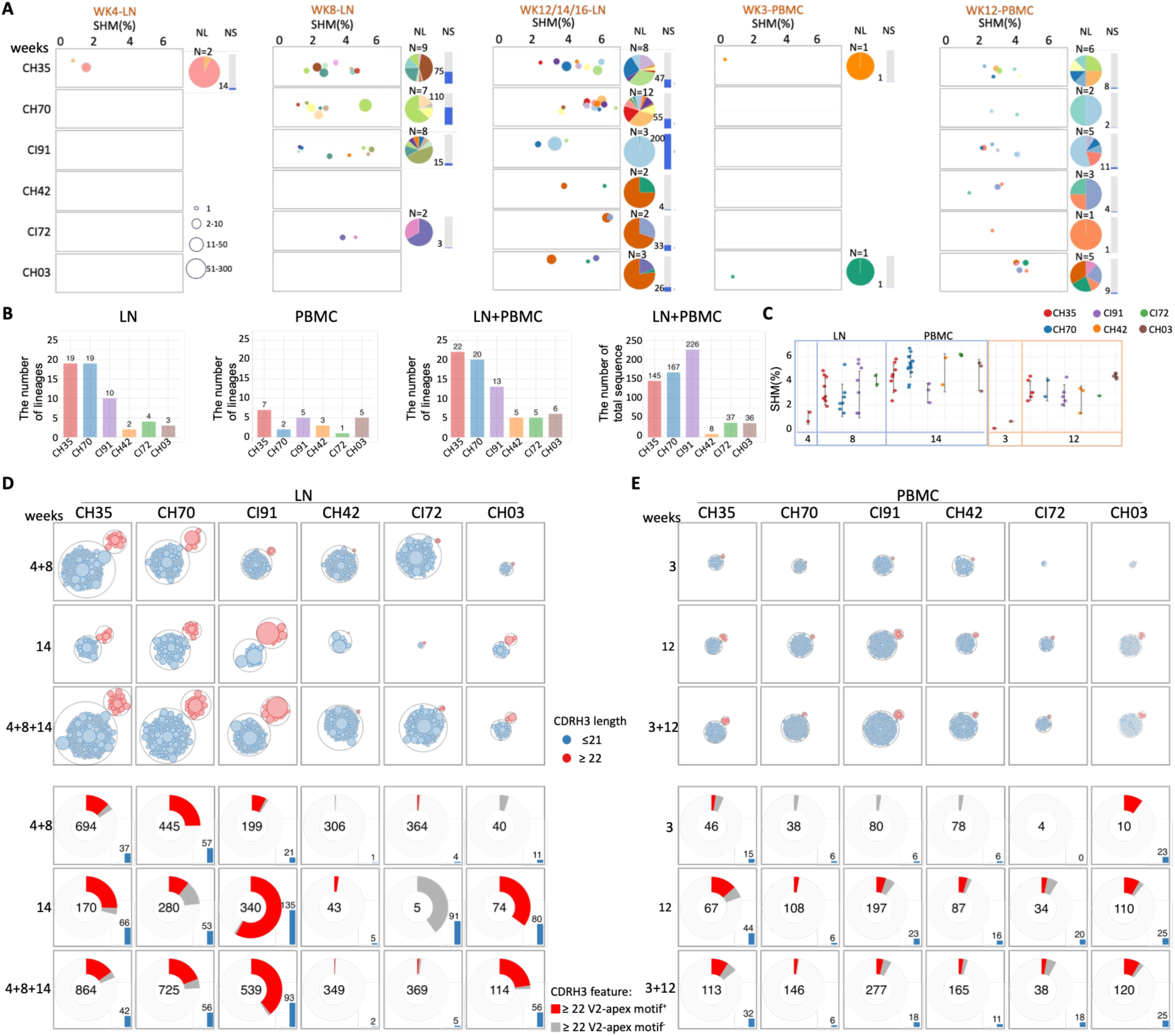
Diversity and composition of epitope-specific V2-apex long CDRH3 loop B cell lineages induced by Q23-APEX-GT2 priming and homologous boosting. (**A**) Bubble plots showing epitope-specific V2-apex Ab-like B cell lineages in LN samples at week 4, week8 and week 12/14/16, and in PBMC samples at week 3 and week 12. Each bubble represents a unique lineage, with size proportional to the number of clonotypes and color distinguishing lineages within each animal. The x-axis denotes somatic hypermutation (SHM) levels, and each row represents one animal. Pie charts summarize the proportion and total number of lineage members per animal at each time point. Bar plots show the total number of V2-apex–like sequence (clonotypes) per animal at the indicated time points. (**B** and **C**) Quantitative summary of isolated V2-apex Ab-like B cell lineages, including: the number of lineages, the total number of sequences and longitudinal SHM levels for each animal. (**D** and **E**) Lineage composition according to CDRH3 length and V2-apex Ab-associated motif usage in epitope-specific IgG⁺ B cells from LNs (D) and PBMCs (E) analyzed by 10x Genomics single-cell sequencing. Circle packing plots (upper) showing lineage distribution according to CDRH3 length. Each bubble represents a unique B cell lineage, with bubble size proportional to the number of clonotype members. Red circles indicate lineages with CDRH3 lengths ≥22 amino acids, whereas blue circles indicate lineages with CDRH3 lengths <22 amino acids. Filled circles denote expanded lineages. Donut charts (bottom) showing the proportion of lineages with CDRH3 lengths ≥22 amino acids that contain conserved V2-apex Ab–associated motifs (red) or lack these motifs (gray) among epitope-specific sorted B cell lineages. Numbers within each donut indicate total sequence counts. Bar plots summarize the fold enrichment of lineages with CDRH3 lengths ≥22 amino acids in immunized rhesus macaques relative to naïve controls. CH35, CH70, and CI91 exhibited a greater proportion of long-CDRH3 lineages containing V2-apex Ab-associated motifs following priming, whereas CH03 showed increased enrichment of V2-apex bnAb-like lineages after homologous boosting.

**fig. S7.**
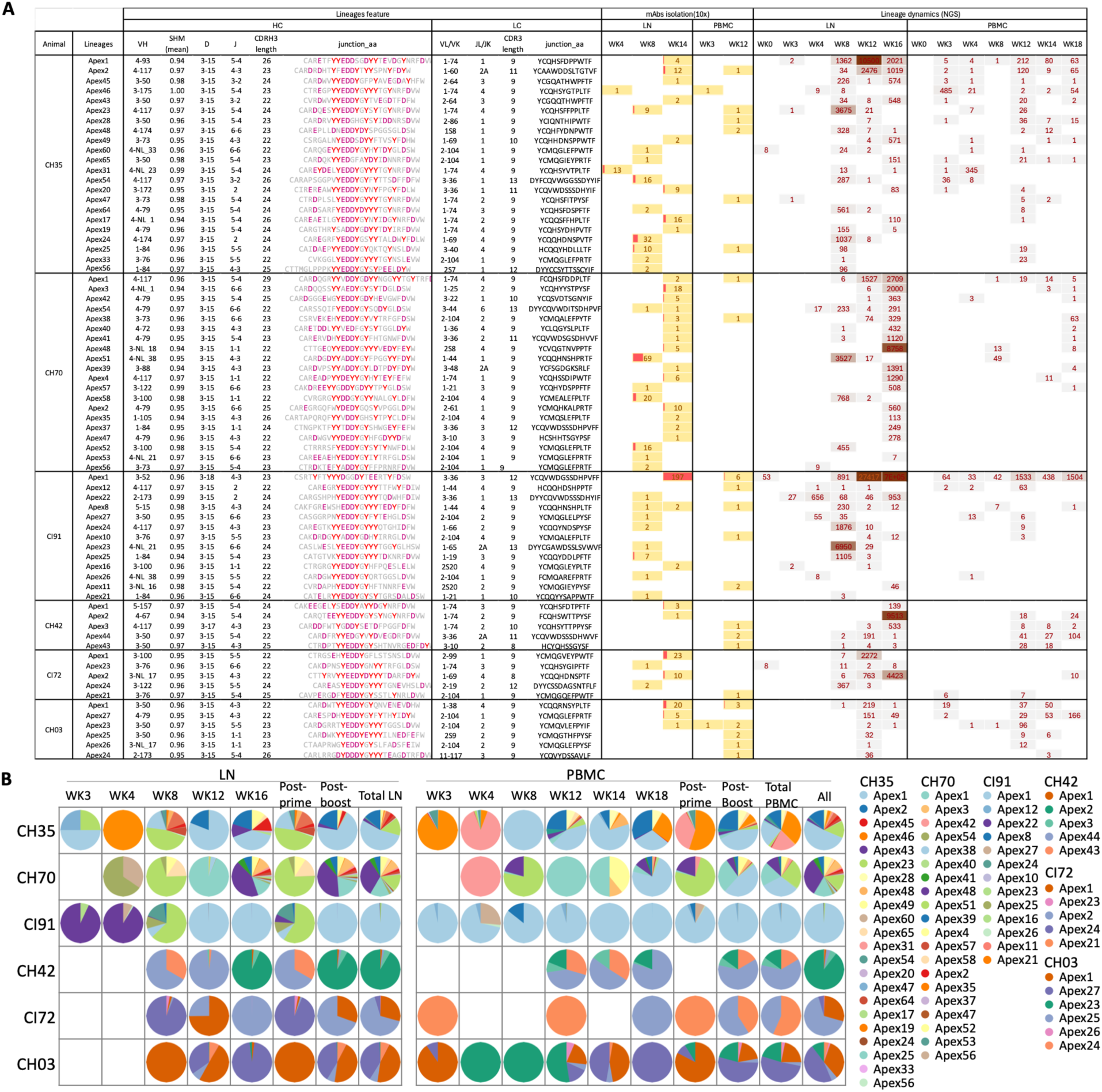
Genetic characteristics and longitudinal tracking of isolated V2-apex Ab-like lineages with CDRH3 lengths ≥22 amino acids across all animals. (**A**) Summary table of all V2-apex Ab-like B cell lineages with CDRH3 lengths ≥22 amino acids isolated via 10x Genomics single-cell sorting from lymph nodes (LNs; weeks 4, 8 and 14) and peripheral blood mononuclear cells (PBMCs; weeks 3 and 12) across all animals. CH42-Apex1 was isolated by week 16 LN single-cell sorting, and CI72-Apex1 and CI72-Apex2 were isolated by week 12 LN single-cell sorting. For each lineage, the table includes: heavy chain (HC) V(D)J gene usage, CDRH3 length, and junction amino acid sequence, light chain (LC) VJ gene usage, CDRL3 length, and junction amino acid sequence, tissue and time point of isolation (e.g., LN-week 4, PBMC-week 12) and longitudinal lineage tracking based on bulk NGS data from LN and PBMC compartments (**B**) Pie charts illustrating lineage distribution in bulk NGS datasets over time in both LN and PBMC compartments. This comprehensive dataset enables cross-sectional and longitudinal comparisons of B cell lineage expansion, persistence, and maturation across compartments and timepoints

**fig. S8.**
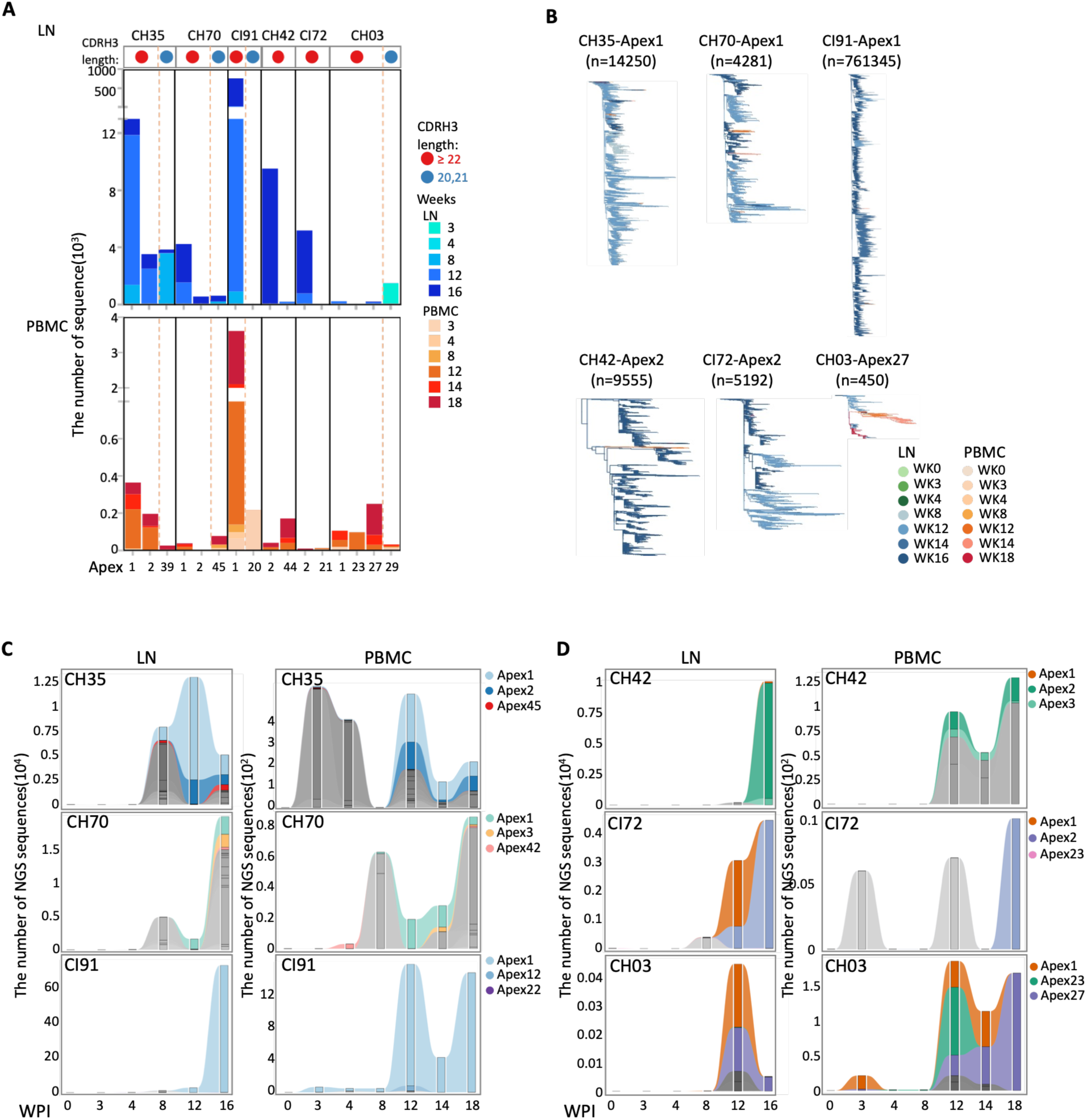
Quantitative tracing of V2-apex Ab-like lineage expansion. (**A**) Bar plots showing the number of sequences for representative lineages with CDRH3 ≥22 amino acids and selected expanded lineages with CDRH3 lengths of 20-21 amino acids, in LN and PBMC. (**B**) Phylogenetic analysis of dominant V2-apex Ab-like lineages. Phylogenetic trees illustrating dominant V2-apex Ab-like B cell lineages isolated from all animals after priming and boosting, corresponding to those shown in Figure 4. Lineages include CH35-Apex1, CH70-Apex1, CI91-Apex1, CH42-Apex2, CI72-Apex2, and CH03-Apex27. Each lineage is phylogenetically clustered with longitudinal B cell sequences derived from LN (weeks 0, 3, 4, 8, 12, 14 and 16) and PBMC (weeks 0, 3, 4, 8, 12, 14, and 18) samples, as determined by bulk NGS. CH35-Apex1, CH70-Apex1 and CI91-Apex1, exhibited extensive clonal diversification and progressive somatic hypermutation. (**C** and **D**) Sankey diagram showing the lineage expansion trends over time for all animals. NGS data from pre-infection time points (weeks 0, 3, 4, 8, 12, and 16 for LN; weeks 0, 3, 4, 8, 12, 14, and 18 for PBMC) were used. Among all long CDRH3 lineages (CDRH3 length ≥22), the top three expanded lineages are color-coded, while all other lineages are shown in grayscale. Overall, dominant lineages in each animal exhibited variable degrees of expansion, with particularly pronounced expansion observed for CH35-Apex1 and CI91-Apex1.

**fig. S9.**
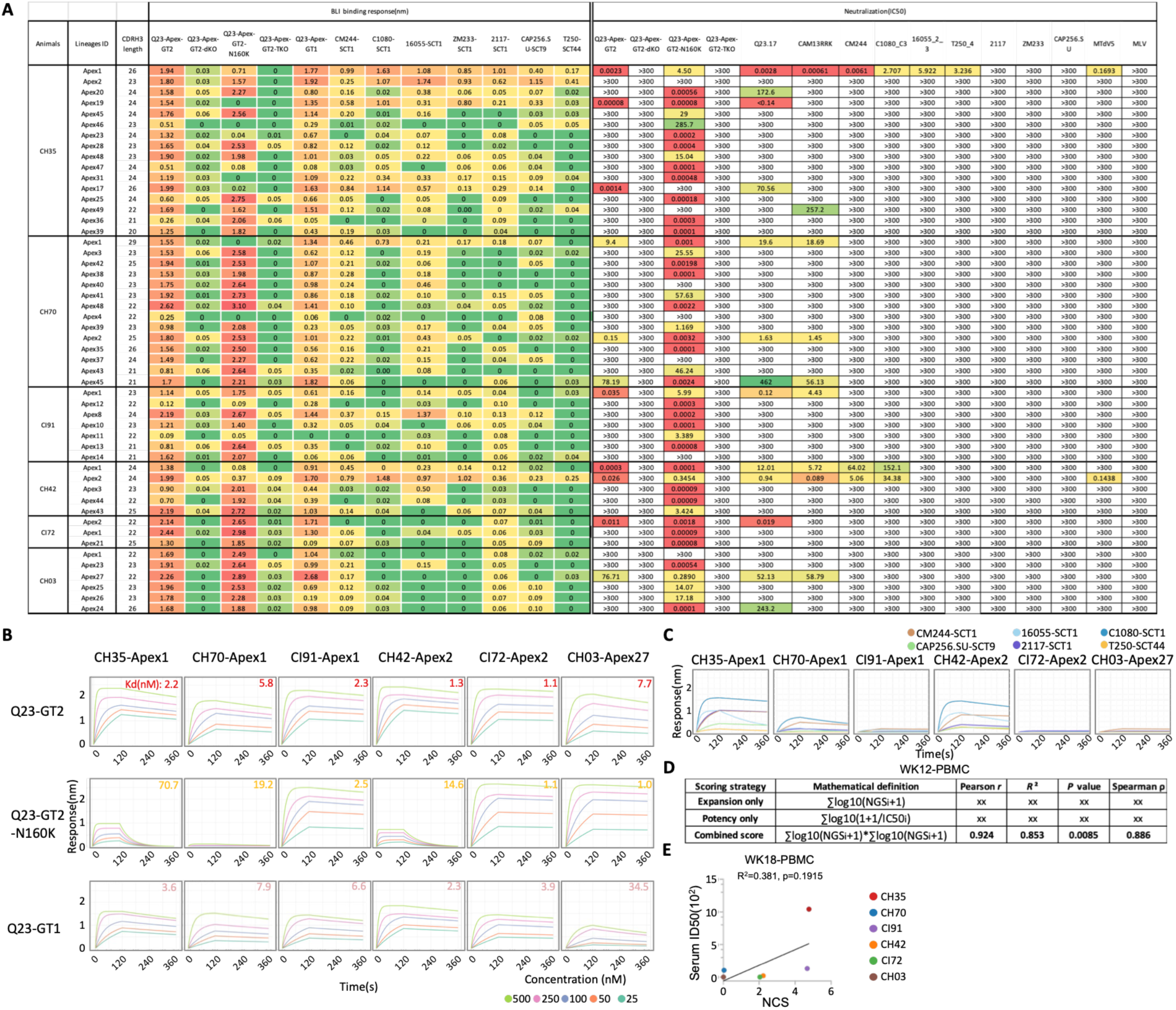
Binding and neutralization features of lineages isolated in Figure 4. (**A**) Maximum BLI binding responses and IC50 neutralization titers are shown for the Q23-APEX-GT2 trimer and its corresponding virus, as well as for the dKO, N160K and N160K-dKO variants, Q23_17 and a panel of V2-sensitive trimers and their corresponding viruses (CM244, C1080_C3, 16055_2_3, ZM233, 2117, CAP256.SU, and T250_4). Neutralization activity against the engineered virus CAM13RRK and the SIV MT145 is also shown. (**B**) BLI titration binding curves of representative lineage Fabs, with Kd indicated in the upper right corner. (**C**) Heterologous trimer binding curves of dominant lineage mAbs from each animal. CH35-Apex1 and CH42-Apex2 exhibited broad heterologous binding and CH70-Apex1 exhibited binding to a subset of heterologous trimers. (**D**) Comparison of biologically motivated scoring strategies for estimating serum neutralization activity. To evaluate the relative contributions of lineage expansion and antibody potency to serum neutralization, three parameter-free scoring strategies were compared using week 12 PBMC lineage abundance and matched serum neutralization ID50. Expansion-only scores considered lineage abundance alone, potency-only scores considered monoclonal antibody neutralization potency alone, and the combined score integrated both variables without parameter fitting. Correlations with serum neutralization were evaluated using Pearson and Spearman analyses. The combined score showed the strongest correlation with serum neutralization and was therefore used for subsequent analyses. (**E**) Scatter plot showing the relationship between the Neutralization Contribution Score (NCS) and serum neutralization ID50 at week 18 PBMC.

**fig. S10.**
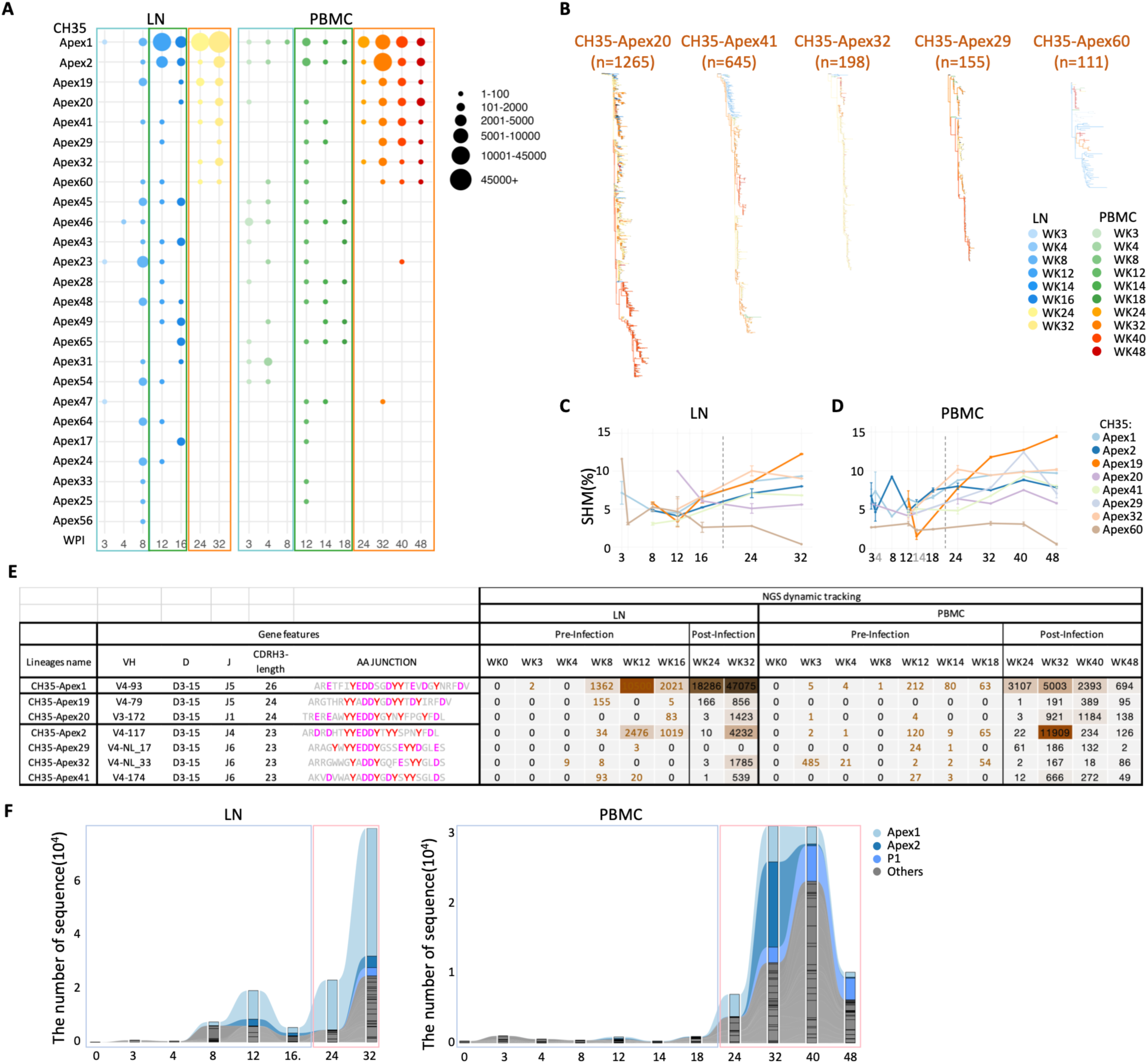
Lineage Tracking and Evolution by Bulk NGS for CH35 recalled lineages. (**A**) Dynamic bubble plots illustrating temporal and compartment-specific changes of all isolated V2-apex lineages across multiple time points, including post-prime (weeks 3, 4 and 8), post-boost (weeks 12, 14, 16 and 18), and post-infection (weeks 24, 32, 40, and 48), in both LNs and PBMCs. Circle size is proportional to the number of clonotypes. Sequences were derived from bulk B cell repertoire sequencing. Several lineages persisted from priming through boosting phases, and some immunogen-elicited lineages were successfully recalled following CAP256.SU SHIV infection. In addition, certain lineages were uniquely detected during the priming, boosting, or post-infection phases. (**B**) Phylogenetic trees illustrating the longitudinal evolution of additional putative recalled lineages identified in fig. S10A. Most lineages were detected in both pre- and post-infection samples and exhibited increased clonal expansion following CAP256.SU infection, consistent with recall and reactivation upon viral exposure. In contrast, Apex60 showed limited post-infection expansion and maturation, suggesting that it is unlikely to represent a bona fide recalled lineage. (**C** and **D**) Longitudinal somatic hypermutation (SHM) trajectories of putative recalled lineages in LNs (C) and PBMCs (D) based on bulk BCR sequencing. Most putative recalled lineages accumulated increasing SHM over time, consistent with continued affinity maturation following infection. In contrast, Apex60 showed minimal SHM accumulation, further supporting that it is unlikely to represent a recalled lineage. (**E**) Table summarizing recalled lineages, including heavy chain (HC) V(D)J gene usage, CDRH3 amino acid sequences, and clonotype counts identified by bulk NGS at pre- and post-infection time points. Darker shading indicates higher clonotype counts and greater lineage expansion. (**F**) Sankey diagram illustrating lineage expansion and dynamics for CH35 over time in LN (left) and PBMC (right) compartments. The most expanded lineages are color-coded, whereas all other lineages are shown in grayscale. Apex lineages refer to those previously confirmed to encode functional V2-apex antibodies. Lineages marked with “P(#)” denote those detected in multiple LN and PBMC NGS samples at both pre- and post-infection time points, with total bulk NGS sequence counts exceeding 1,000. Notably, CH35-Apex1 exhibited sustained clonal expansion, with a marked increase in lineage abundance following CAP256.SU infection.

**fig. S11.**
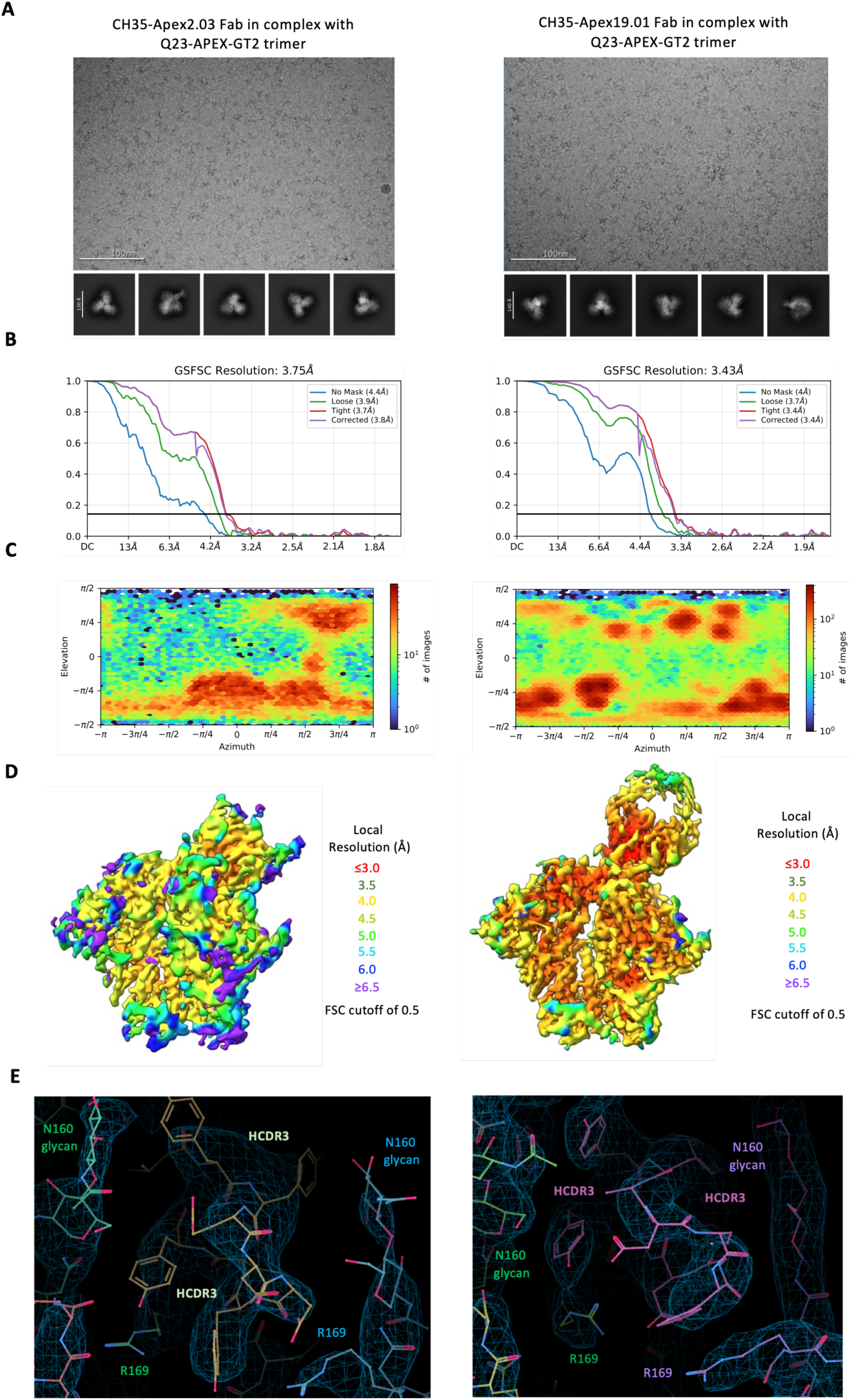
Single-particle cryo-EM validation for SHIV.CAP256SU-recalled V2 apex lineages in complex with Q23-APEX-GT2 envelope trimer. (**A**) Representative raw micrograph with representative 2D class averages of picked particles shown below. (**B**) Orientations of all particles used in the final refinement are shown as a heatmap. (**C**) Gold-standard fourier shell correlation (FSC) curves with auto-tightening using a non-uniform refinement with C1 symmetry. (**D**) Local resolution estimation of the full map is shown as generated through cryoSPARC using an FSC cutoff of 0.5. (**E**) Example of high-resolution cryo-EM 3D reconstruction density to highlight the Fab HCDR3 interface with N-linked glycans and R169 residues.

**fig. S12.**
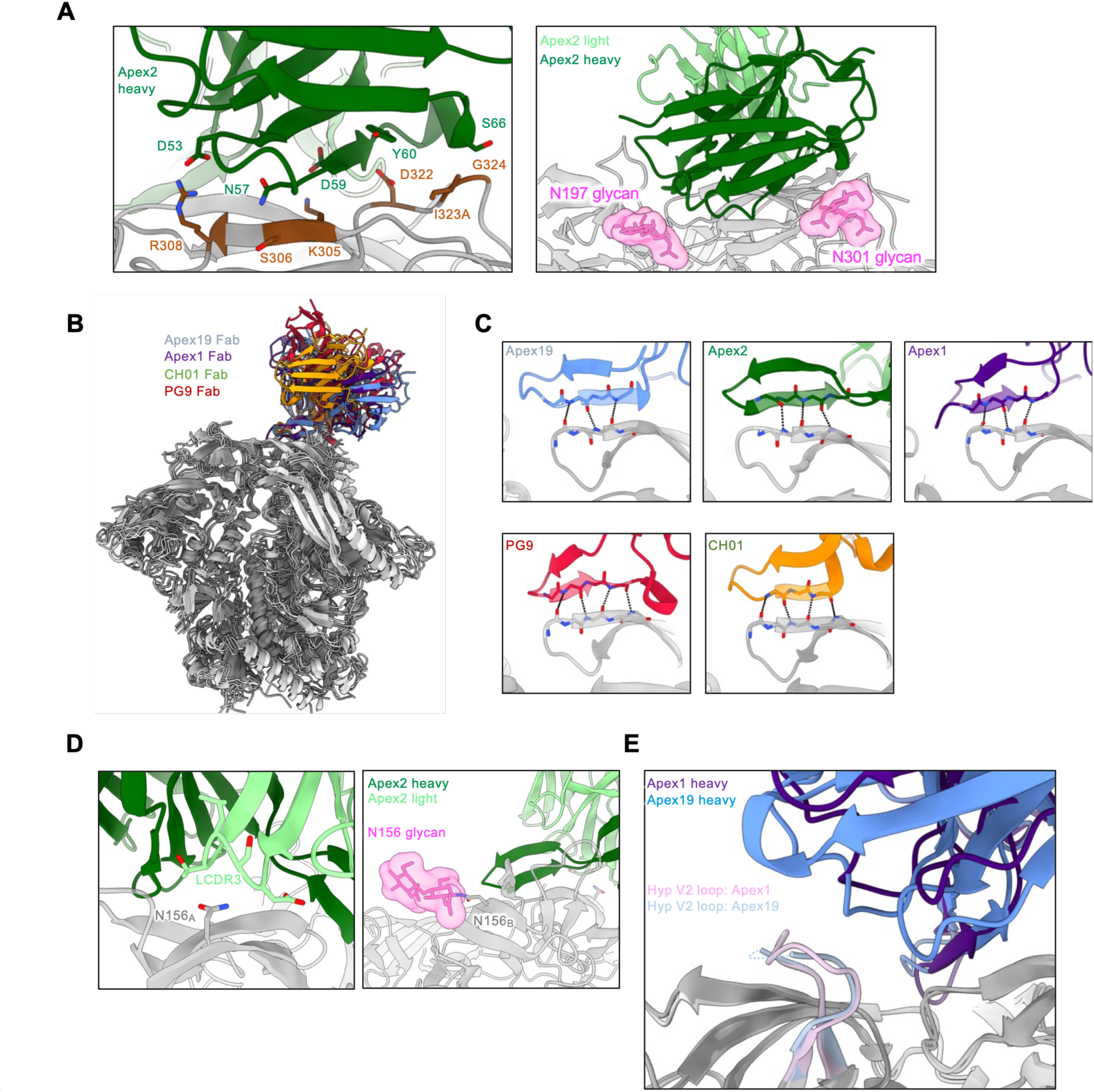
Structural analysis of SHIV.CAP256SU-recalled lineages. (**A**) Expanded interface views of CH35-Apex2.03 recognizing the V3 loop (left) and non-V2 apex epitope glycans (right). (**B**) Structural alignment of the Fab:trimer complexes of rhesus antibodies CH35-Apex1.08 and CH35-Apex2.03 with human antibodies PG9 (PDB-7t77) and CH01 (PDB-9dhw). (**C**) Rhesus antibodies CH35-Apex1.08, Apex2.03, and Apex19.01 (top) and human antibodies PG9 and CH01 (bottom) parallel main-chain bonding with the C-strand via HCDR3. Hydrogen bonds are depicted with dotted lines. (**D**) Env N156 residues on the primary protomer A (left) and adjacent protomer B (right) from the Apex2.03 complex. (**E**) Structural alignment of CH35-Apex1.08 and Apex19.01 to compare recognition of the hypervariable V2 loop from adjacent protomer B. Three V2 loop residues that are disordered in the Apex19.01 structure are indicated with a dashed line.

**fig. S13.**
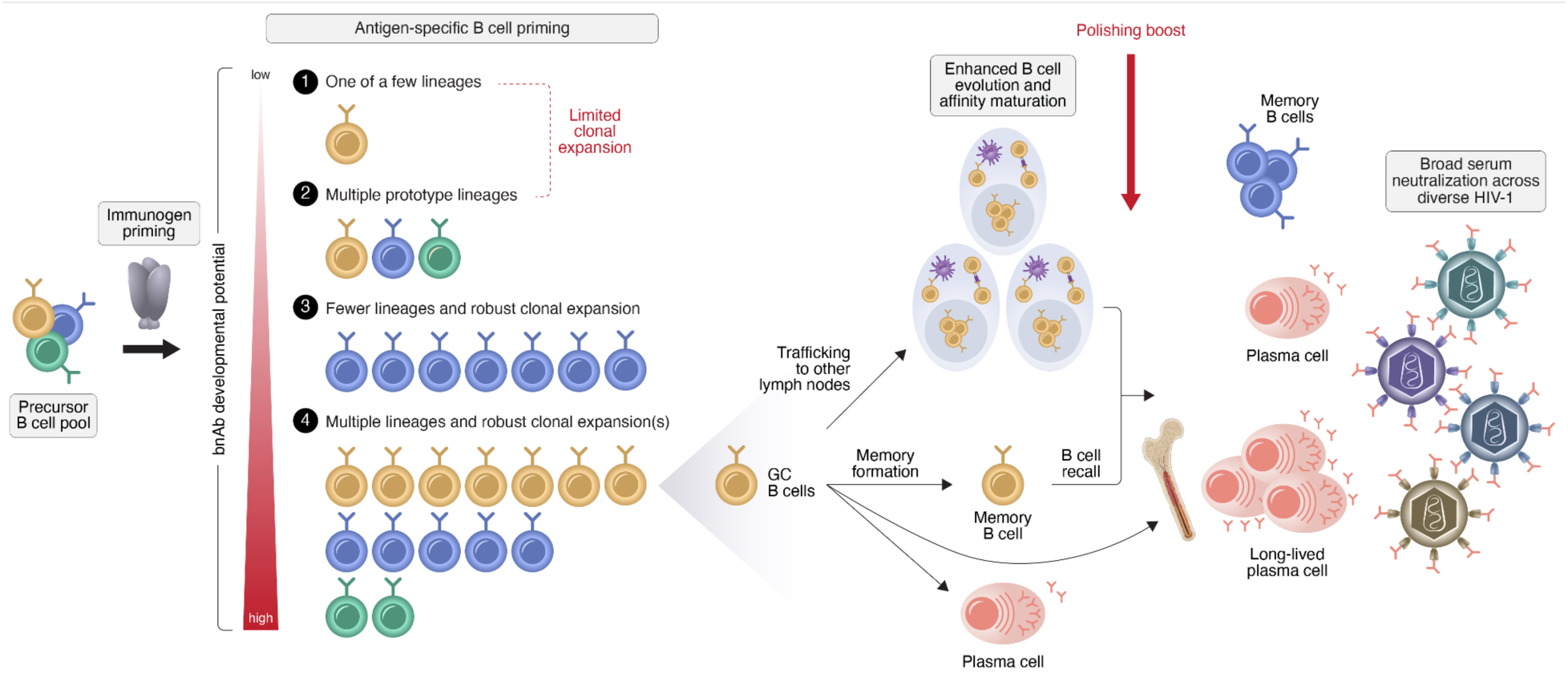
Model of efficient rare bnAb B cell priming and early clonal dominance driving serum neutralization. Broad precursor recruitment and robust early clonal expansion are key determinants of germline-targeting immunogen priming efficiency. A germline-targeting immunogen that efficiently primes multiple diverse bnAb precursor lineages and robustly expands a subset of these lineages is likely to have the greatest potential for generating serum broadly neutralizing antibody responses. Dominant clonal lineages that expand early can traffic across lymphoid sites, enhancing opportunities for continued evolution, re-encounter with boosting immunogens, and further affinity maturation. These lineages may differentiate into memory B cells—preserving recall potential— or plasma cells that directly contribute to serum neutralization. Upon boosting, germinal center and memory B cells are reactivated, undergoing additional expansion and selection that amplifies plasma cell output and antibody production. A subset of recalled cells is retained as memory, sustaining responsiveness to future antigen exposure. Some may be deposited in bone marrow as long-lived plasma cells, supporting durable antibody secretion and long-lasting serum neutralization.

**Table S1. Longitudinal analysis of V2-apex epitope–specific serum antibody responses following Q23-APEX-GT2 immunization.**

Table summarizing ID_50_ neutralization titers measured in longitudinal serum samples from rhesus macaques immunized with the Q23-APEX-GT2 trimer. Sera were evaluated against the homologous viruses Q23-APEX-GT2 and Q23_17, Q23-APEX-GT2 mutant variants (dKO, N160K, and N160K-dKO), and a panel of V2-apex–sensitive heterologous viruses, including CAM13RRK, CM244, CAP256.SU, V703_2117, ZM233_6, and 25710_2_43.

**Table S2. Biolayer interferometry (BLI) binding analysis of CH35 recalled mAb lineages**. Table summarizing maximum BLI binding responses for V2-apex–directed monoclonal antibody (mAb) lineages isolated from animal CH35 after Q23-APEX-GT2 trimer immunization and following CAP256.SU SHIV infection. Comparative analysis of pre- and post-infection mAbs demonstrates that lineages recovered after infection exhibit increased binding magnitude and enhanced cross-reactivity to a panel of heterologous Env trimers, consistent with further affinity maturation and expansion of breadth.

**Table S3. Neutralization IC50 profiles of CH35 recalled mAb lineages.** Table summarizing neutralization IC50 values for CH35 V2-apex–directed monoclonal antibody (mAb) lineages isolated before and after CAP256.SU SHIV infection. Comparative analysis of pre- and post-infection antibodies reveals that CH35-Apex1 and CH35-Apex19 acquired increased neutralization breadth and potency following infection, consistent with enhanced functional maturation. In contrast, CH35-Apex2 displayed expanded binding breadth but lacked detectable neutralization activity both before and after infection, indicating that increased Env reactivity did not translate into functional neutralization capacity for this lineage.

**Table S4. Data acquisition, processing details and final model statistics for all structures.**

**Table S5. mAb panel used for flow cytometry staining.**

## Notes

### Summary of Updates

The manuscript has been substantially revised and restructured to improve the overall narrative, strengthen the presentation of the findings, and better highlight the significance of the study. We have also incorporated new data that provide additional support for our key findings and strengthen the conclusions of the paper.

